# Cortical structure in relation to empathy and psychopathy in 800 incarcerated men

**DOI:** 10.1101/2023.06.14.543399

**Authors:** Marcin A. Radecki, J. Michael Maurer, Keith A. Harenski, David D. Stephenson, Erika Sampaolo, Giada Lettieri, Giacomo Handjaras, Emiliano Ricciardi, Samantha N. Rodriguez, Craig S. Neumann, Carla L. Harenski, Sara Palumbo, Silvia Pellegrini, Jean Decety, Pietro Pietrini, Kent A. Kiehl, Luca Cecchetti

## Abstract

**Background:** Reduced empathy is a hallmark of individuals with high (i.e., clinical) levels of psychopathy, who are overrepresented among incarcerated men. Yet, a comprehensive, well-powered mapping of cortical structure in relation to empathy and psychopathy is still lacking.

**Methods:** In 804 incarcerated adult men, we administered the Perspective Taking (IRI-PT) and Empathic Concern (IRI-EC) subscales of the Interpersonal Reactivity Index, the Psychopathy Checklist-Revised (PCL-R; with Interpersonal/Affective [F1] and Lifestyle/Antisocial [F2] factors), and T1-weighted MRI to quantify cortical thickness (CT), surface area (SA), and structural-covariance gradients.

**Results:** PCL-R F1 was uniquely negatively related to IRI-EC, while PCL-R F2 was uniquely negatively related to IRI-PT. Cortical structure was not related to the IRI subscales. In contrast, CT was related to PCL-R F1 (mostly positively), SA was related to both PCL-R factors (only positively), and both cortical indices demonstrated out-of-sample predictive utility for PCL-R F1. Compared to low-psychopathy men, high-psychopathy men had uniquely lower IRI-EC scores and increased SA (but not CT); across the cortex, effect sizes were largest in the paralimbic class and somatomotor network, while meta-analytic task-based activations further highlighted spatial overlap with (social-)affective/sensory regions. Finally, the total sample revealed canonical anterior-posterior structural-covariance gradients. In high-psychopathy men, the gradient of CT (but not SA) was globally compressed.

**Conclusions:** High-psychopathy men had reduced empathic concern, increased SA, and a compressed macroscale organization of CT, suggesting specific co-occurring alterations in empathy and cortical structure. Future work should build upon these novel insights in both the general and incarcerated populations to inform the treatment of psychopathy.

## Introduction

Empathy comprises desirable psychological traits that allow us to understand another person’s mental states (cognitive empathy), share their affective states (affective empathy), and feel concern for their well-being (empathic concern) (1–6). Reduced empathy – particularly affective empathy and empathic concern – is a hallmark of individuals with high (i.e., clinical) levels of psychopathy (7-9; for meta-analyses, see (10,11)). More broadly, psychopathy is a constellation of interpersonal/affective traits (e.g., said reduced empathy) and lifestyle/antisocial traits (e.g., impulsivity), as operationalized by Factors 1 and 2 of the Psychopathy Checklist-Revised (PCL-R), respectively (13–14). The prevalence of high psychopathy based on the PCL-R approximates 1.2% in the general population; it is higher among incarcerated individuals (up to 25%) and among males/men than females/women (up to three-fold) (15). Because psychopathy incurs societal costs that reach hundreds of billions of USD per annum through violence, crime, and recidivism (16), we need to advance its phenotypic and neuroimaging characterization to improve long-term treatment outcomes (17–19).

It remains unknown how brain structure is related to empathy (e.g., as measured by the Interpersonal Reactivity Index; IRI (20–23)) in the incarcerated population. Even in the general population, no structural meta-analysis has yet been conducted, further motivating the investigation into the underlying brain structure (for study examples, see 23-25)). Functional meta-analyses highlight default-mode hubs for cognitive empathy (e.g., medial prefrontal cortex), ventral-attention hubs for affective empathy (e.g., insula), and reward-related regions for empathic concern (e.g., striatum and ventromedial prefrontal cortex) (26–31). Altered function of these and more distributed regions, particularly during social and affective processing (32–40), provides insights into reduced empathy and other psychopathic traits (for meta-analyses, see (41–43)). While the structural literature on psychopathy is extant (44–49), it suffers from limited replicability, in part owing to small sample sizes (50), and is almost exclusively focused on voxel-based gray-matter volume (GMV), with the most common result being reduced GMV (for a meta-analysis, see (51); see also 52)). Surprisingly, no study on psychopathy to date has investigated (raw) cortical surface area (SA), let alone alongside cortical thickness (CT), the two critical constituents of cortical GMV (but see (53)). Given that CT and SA exhibit distinct evolutionary (54), genetic (55), developmental (56), and psychiatric (57) profiles, it is imperative to fractionate them, as they may differentially map onto multidimensional empathy and psychopathy. Indeed, SA has already been demonstrated to have a higher sensitivity than CT to broadly construed antisocial behavior (58,59; see also (60)). To further enhance the generalizability of such brain-behavior relationships, a multivariate framework with out-of-sample predictive modeling is recommended (61–63), particularly given that CT and SA have not yet demonstrated predictive utility for psychopathy (but see (64)).

To make this cortical mapping even more comprehensive, and to overcome some of the limitations of investigating raw cortical structure, a novel framework is offered by gradients: topographical patterns of regional similarity, including structural covariance (65–67), embedded in a low-dimensional space (68–70). Based on meta-analytic data, CT gradients have been shown to differ across major psychiatric conditions in a transdiagnostic fashion (71–73) that dovetails with differences along the primary (i.e., unimodal-transmodal) axis of intrinsic connectivity in schizophrenia (74), autism (75), and depression (76). In these conditions, the unimodal-transmodal gradient has been observed to be compressed (as opposed to expanded (77)). This compression corresponds to a smaller gradient range and suggests reduced differentiation between its ends, where the opposing unimodal/sensorimotor regions and transmodal/association regions have more similar connectivity patterns (78). Importantly, an anterior-posterior compression has also been observed for a CT gradient in schizophrenia (79). It remains to be established whether high-psychopathy individuals may exhibit similar differences in the macroscale organization of either CT or SA.

In sum, a comprehensive, well-powered mapping of cortical structure in relation to empathy and psychopathy is still lacking. There is thus a need to investigate CT alongside SA in an interpretable way that is facilitated by theories of psychopathy (e.g., regarding laminar differentiation (80–82)) and brain function (e.g., regarding intrinsic connectivity (83)). Simultaneously, this gap presents an opportunity to clarify statistically unique relationships between the IRI and PCL-R, two of the most widely used measures of empathy and psychopathy, respectively (12). Here, we address these gaps in a large sample of incarcerated adult men (N = 804), grounding our expectations in meta-analyses documenting negative relationships of psychopathy with both empathy (10,11) and cortical GMV (51). We ask five overarching questions (see also *Statistical analysis*):

Q1: How is psychopathy related to empathy given the multidimensionality of both constructs and their potential for unique relationships?
Q2: How is cortical structure, distinguishing CT and SA, related to empathy and psychopathy?
Q3: Can cortical structure predict empathy and psychopathy in out-of-sample individuals?
Q4: How does cortical structure differ among individuals with high (i.e., clinical) levels of psychopathy?
Finally, Q5: How do structural-covariance gradients differ among these individuals?

## Methods and Materials

### Participants

Nine-hundred-twelve adult men (gender self-reported) were recruited by the Mind Research Network (MRN) from correctional facilities in the southwestern and midwestern United States, and had partial data available, including a T1-weighted MRI scan. We included N = 804 who sequentially met the following criteria: (1) passed MRI quality control (N_excluded_ = 105); (2) had data on empathy, psychopathy, age, and IQ (N_excluded_ = 1); and (3) had an IQ ≥ 70 (N_excluded_ = 2) (for participant characteristics, see *Table **1***). Among the included participants, N = 723 (∼90%) reported having committed a violent crime (e.g., murder), N = 715 (∼89%) a non-violent crime (e.g., theft), and N = 634 (∼79%) both a violent and non-violent crime (based on a classification similar to (84); see *Covariates* in the Supplement). All participants provided written informed consent, and all research procedures were approved by the Institutional Review Board of the University of New Mexico or the Ethical and Independent Review Services for data collection post June 2015.

**Table 1.**
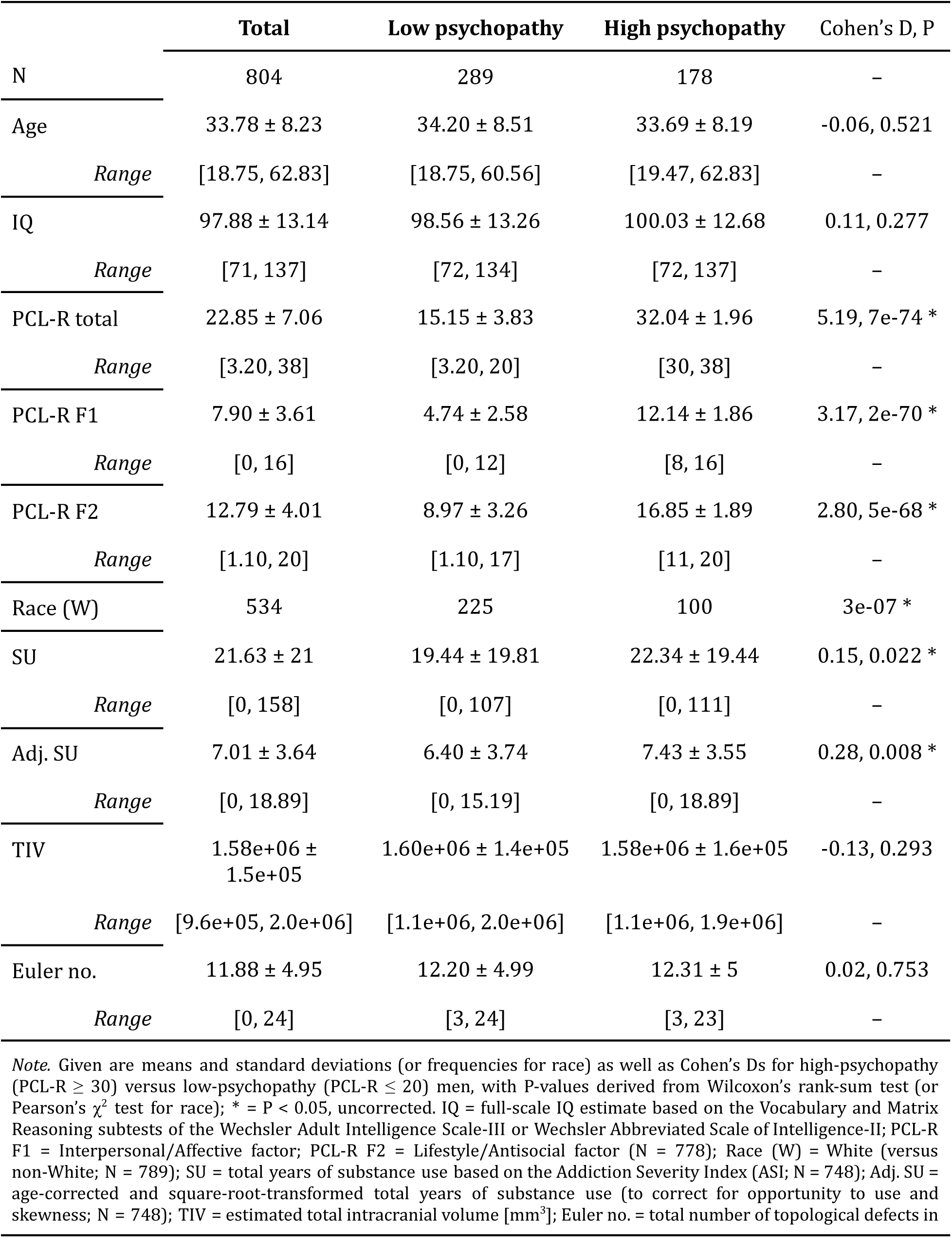
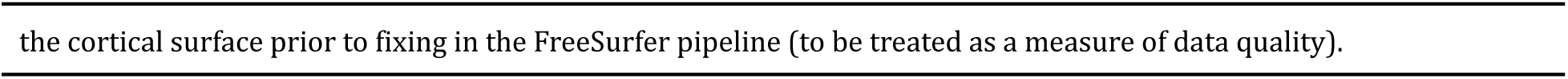
Participant characteristics.

|  | **Total** | **Low psychopathy** | **High psychopathy** | **Cohen's D, P** |
| --- | --- | --- | --- | --- |
| N | 804 | 289 | 178 | – |
| Age | 33.78 ± 8.23 | 34.20 ± 8.51 | 33.69 ± 8.19 | -0.06, 0.521 |
| *Range* | [18.75, 62.83] | [18.75, 60.56] | [19.47, 62.83] | – |
| IQ | 97.88 ± 13.14 | 98.56 ± 13.26 | 100.03 ± 12.68 | 0.11, 0.277 |
| *Range* | [71, 137] | [72, 134] | [72, 137] | – |
| PCL-R total | 22.85 ± 7.06 | 15.15 ± 3.83 | 32.04 ± 1.96 | 5.19, 7e-74 * |
| *Range* | [3.20, 38] | [3.20, 20] | [30, 38] | – |
| PCL-R F1 | 7.90 ± 3.61 | 4.74 ± 2.58 | 12.14 ± 1.86 | 3.17, 2e-70 * |
| *Range* | [0, 16] | [0, 12] | [8, 16] | – |
| PCL-R F2 | 12.79 ± 4.01 | 8.97 ± 3.26 | 16.85 ± 1.89 | 2.80, 5e-68 * |
| *Range* | [1.10, 20] | [1.10, 17] | [11, 20] | – |
| Race (W) | 534 | 225 | 100 | 3e-07 * |
| SU | 21.63 ± 21 | 19.44 ± 19.81 | 22.34 ± 19.44 | 0.15, 0.022 * |
| *Range* | [0, 158] | [0, 107] | [0, 111] | – |
| Adj. SU | 7.01 ± 3.64 | 6.40 ± 3.74 | 7.43 ± 3.55 | 0.28, 0.008 * |
| *Range* | [0, 18.89] | [0, 15.19] | [0, 18.89] | – |
| TIV | 1.58e+06 ± 1.5e+05 | 1.60e+06 ± 1.4e+05 | 1.58e+06 ± 1.6e+05 | -0.13, 0.293 |
| *Range* | [9.6e+05, 2.0e+06] | [1.1e+06, 2.0e+06] | [1.1e+06, 1.9e+06] | – |
| Euler no. | 11.88 ± 4.95 | 12.20 ± 4.99 | 12.31 ± 5 | 0.02, 0.753 |
| *Range* | [0, 24] | [3, 24] | [3, 23] | – |

*Note.* Given are means and standard deviations (or frequencies for race) as well as Cohen's Ds for high-psychopathy (PCL-R ≥ 30) versus low-psychopathy (PCL-R ≤ 20) men, with P-values derived from Wilcoxon's rank-sum test (or Pearson's $\chi^2$ test for race); \* = P < 0.05, uncorrected. IQ = full-scale IQ estimate based on the Vocabulary and Matrix Reasoning subtests of the Wechsler Adult Intelligence Scale-III or Wechsler Abbreviated Scale of Intelligence-II; PCL-R F1 = Interpersonal/Affective factor; PCL-R F2 = Lifestyle/Antisocial factor (N = 778); Race (W) = White (versus non-White; N = 789); SU = total years of substance use based on the Addiction Severity Index (ASI; N = 748); Adj. SU = age-corrected and square-root-transformed total years of substance use (to correct for opportunity to use and skewness; N = 748); TIV = estimated total intracranial volume [mm<sup>3</sup>]; Euler no. = total number of topological defects in
the cortical surface prior to fixing in the FreeSurfer pipeline (to be treated as a measure of data quality).

We also included the male sample from the Human Connectome Project (HCP) Young Adult S1200 release (85–87) with structural-MRI and IQ data (N = 501; *Supplementary Table **S1***). This dataset has been used to derive the canonical anterior-posterior gradient of CT (67), which then served as an external reference in the aforementioned study on schizophrenia (79). Given that structural-covariance gradients in our total sample served as the alignment reference for psychopathy groups (Q5), it was important to evaluate their normativity (i.e., consistency with the general male population, represented by the HCP sample), and for a related sensitivity analysis by psychopathy group. Note that the HCP did not include our measures of empathy and psychopathy.

### MRI

On the grounds of the correctional facilities, high-resolution T1-weighted MRI scans were acquired with the MRN’s mobile scanner (i.e., 1.5-T Siemens MAGNETOM Avanto with a 12-channel, multi-echo MPRAGE pulse sequence). The scanning parameters were as follows: repetition time = 2,530 ms; echo times = 1.64 ms, 3.50 ms, 5.36 ms, and 7.22 ms; inversion time = 1,100 ms; flip angle = 7°; slice thickness = 1.3 mm; matrix size = 256 × 256, yielding 128 sagittal slices with an in-plane resolution of 1.0 mm × 1.0 mm.

Each scan underwent the standard recon-all pipeline in FreeSurfer version 7.4.1 (https://surfer.nmr.mgh.harvard.edu/ (88)) and was parcellated in the HCP-MMP1.0 atlas to delineate 360 regions (89). For quality control and HCP data, see *MRI* in the Supplement.

### Empathy

Empathy was measured with the Perspective Taking (IRI-PT) and Empathic Concern (IRI-EC) subscales of the Interpersonal Reactivity Index (IRI) (21), a self-report questionnaire of trait empathy widely used in both community and incarcerated samples. Across these samples, IRI-PT is traditionally labeled a measure of cognitive empathy while IRI-EC a measure of affective empathy (10,11,90–93). While we agree with the labelling of IRI-PT, the labelling of IRI-EC conflates affective empathy with empathic concern; we label IRI-EC a measure of the latter. More specifically, IRI-PT assesses the “tendency to spontaneously adopt the psychological point of view of others” while IRI-EC the “feelings of sympathy and concern for unfortunate others” ((23), pp. 113–114). Each subscale includes seven items scored on a five-point Likert scale ranging from “Does not describe me well” (0 points) to “Describes me very well” (4 points) (for all items, see *Supplementary Table **S2***). Possible scores thus range 0-28 points per subscale, with higher scores indicating higher empathy. The remaining Fantasy and Personal Distress subscales of the IRI were not included, as they are less frequently used to distinguish cognitive empathy from affective empathy and empathic concern (92), and are less related to psychopathy (10). See also *Internal consistency* in the Supplement.

### Psychopathy

Psychopathy was measured with Hare’s Psychopathy Checklist-Revised (PCL-R) (13). All PCL-R scores were based on both a semi-structured interview and institutional-file review conducted by the MRN research staff with a bachelor’s degree or higher following rigorous training designed and supervised by K.A.K. MRN has historically completed independent double-ratings on ∼10% of all PCL-R interviews, obtaining excellent rater agreement (45). The PCL-R includes 20 items that correspond to two factors: Interpersonal/Affective (F1) and Lifestyle/Antisocial (F2) (for all items, see *Supplementary Table **S3***). Each item is scored 0, 1, or 2 points, indicating no evidence, some evidence, and pervasive evidence, respectively. The total score is a sum across the 20 items, thus ranging 0-40 points, with a higher score indicating higher psychopathy. PCL-R total and PCL-R F1 were available for the total sample (i.e., N = 804, where N = 582 and N = 798 had complete item-level data, respectively); PCL-R F2 was available for N = 778 (where N = 617 had complete item-level data). For items omitted due to insufficient information, we used a prorating formula to estimate the total and factor scores with possible decimals. See also *Internal consistency* in the Supplement.

Following both the PCL-R guideline (14) and extensive work with incarcerated adult males/men (e.g., (32–36,40,94–96)), we defined “high” psychopathy at PCL-R ≥ 30 and “low” psychopathy at PCL-R ≤ 20. Beyond comparability with the literature, this extreme-groups approach allowed us to specifically investigate participants with clinical levels of psychopathy against a clearly isolated non-clinical group, knowing that those highest in psychopathy are at the highest risk for future antisocial behavior (97–99). Further, this approach allowed for comparability with inherently categorical analyses (Q5) while not necessarily reducing statistical power (100). Therefore, we focus on these categorical analyses below (alongside PCL-R F1 and PCL-R F2), but for completeness, report analyses for PCL-R total in the Supplement.

### Statistical analysis

Statistical analysis was run in MATLAB version R2020b (The MathWorks Inc., Natick, MA; https://www.mathworks.com/) and is described in further detail in the Supplement (alongside *Mesulam’s classes and Yeo’s networks*; *Multivariate prediction*; *Meta-analytic task-based activations*; and *Structural-covariance gradients*). Briefly, for Q1, we tested for relationships between psychopathy (PCL-R F1 and PCL-R F2; PCL-R total in the Supplement) with empathy (IRI-PT and IRI-EC), controlling for age and IQ in a robust linear regression. We then tested IRI-PT and IRI-EC by psychopathy group, controlling for the same covariates. For Q2, we tested for relationships of cortical structure (CT and SA) with empathy and psychopathy at a false-discovery rate of P < 0.05 (101), controlling for age and IQ with CT and additionally for total intracranial volume (TIV) with SA. For interpretability, we compared effect sizes across the cortex by four Mesulam’s classes (82) and seven Yeo’s networks (83) using Wilcoxon’s rank-sum test. For Q3, which complemented the univariate analyses for Q2, we leveraged a multivariate framework with a train-test split and cross-validation to predict empathy and psychopathy from cortical structure (corrected for the same covariates) using ridge regression. For Q4, we tested for global and regional differences in cortical structure by psychopathy group, controlling for the same covariates. We then compared effect sizes across the cortex as above; for additional psychological interpretability, we computed spatial overlap between results at P_FDR_ < 0.05 and meta-analytic task-based activations from Schurz et al. (31) and Neurosynth (102). Finally, for Q5, we tested for psychopathy-group differences in CT and SA gradients using Kolmogorov-Smirnov’s and Wilcoxon’s signed-rank tests. Here, gradient consistency between the total and HCP samples was evaluated via spatial correlation (103,104). Across the analyses, Bonferroni’s correction was applied where appropriate.

## Results

### Empathy and psychopathy (Q1)

In 804 incarcerated adult men, we first tested for relationships between empathy and psychopathy (Q1). PCL-R F1 had a negative relationship with IRI-EC, while PCL-R F2 had a negative relationship with both IRI subscales (*Fig. **1A-B***, *Supplementary Fig. **S1***, and *Supplementary Table **S4***). In categorical analyses, high-psychopathy men, compared to low-psychopathy men, scored lower on both IRI subscales, with a larger effect size for IRI-EC. These relationships clarified when additionally controlling for the other IRI subscale: PCL-R F1 was uniquely negatively related to IRI-EC, while PCL-R F2 was uniquely negatively related to IRI-PT. Further, the group difference on IRI-PT became insignificant, while the group difference on IRI-EC remained significant, and this did not change when additionally controlling for race and substance use (*Fig. **1C***, *Supplementary Fig. **S1***, and *Supplementary Table **S5***). For sample-specific correlations, see *Fig. **1D***.

**Figure 1.**
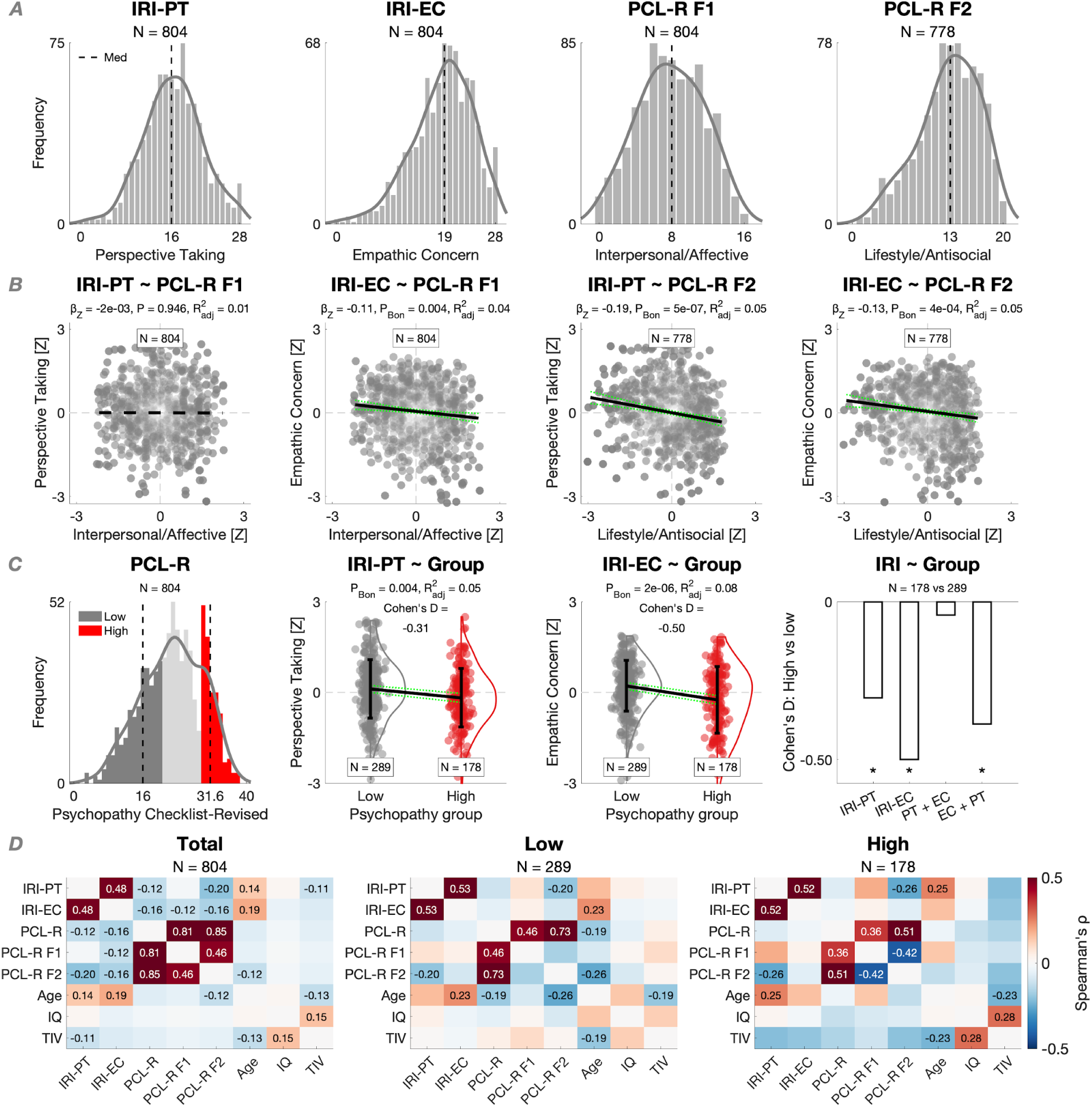
Psychopathy in relation to empathy (Q1). (A) Distribution of IRI-PT, IRI-EC, PCL-R F1, and PCL-R F2. (B) Negative relationships of PCL-R F1 and PCL-R F2 with IRI-PT and IRI-EC, controlling for age and IQ in a robust linear regression, with Bonferroni’s correction across the IRI subscales. (C) From left to right: Distribution of PCL-R total, depicting low-psychopathy (PCL-R ≤ 20; dark gray) and high-psychopathy (PCL-R ≥ 30; red) men; lower scores on IRI-PT and IRI-EC in high-psychopathy men, controlling for age and IQ, with Bonferroni’s correction across the IRI subscales; lower score in high-psychopathy men on IRI-EC but not IRI-PT when additionally controlling for the other IRI subscale. (D) Sample-specific Spearman’s correlation matrices, with numeric effect sizes displayed at P_Bon_ < 0.05 following correction across the 28 tests.

### Cortical structure, empathy, and psychopathy (Q2-Q3)

Next, we tested for relationships of CT and SA with empathy and psychopathy (Q2). CT was not related to the IRI subscales or PCL-R F2. However, CT in 16 parcels had a positive relationship with PCL-R F1, while CT in six parcels had a negative relationship. Across the cortex, effect sizes for PCL-R F1 were largest in the heteromodal class of Mesulam and frontoparietal network of Yeo (positive median in both cases), with differentiation by both class (e.g., heteromodal > paralimbic) and network (e.g., frontoparietal > visual) (*Supplementary Figs. **2***-***3*** and *Supplementary Table **S6***). Similarly to CT, SA was not related to the IRI subscales. In contrast, SA had a positive relationship with PCL-R F1 in 103 parcels and with PCL-R F2 in the same three parcels in the right superior-temporal/auditory cortex. For both PCL-R factors, effect sizes were largest in the paralimbic class and somatomotor network (positive median in both cases), with differentiation by class for PCL-R F2 (e.g., paralimbic > heteromodal) and by network for both PCL-R factors (e.g., somatomotor > dorsal-attention) (*Fig. **2*** and *Supplementary Tables **S7-S9***). In sensitivity analyses, the null CT and SA results for the IRI subscales did not change when leveraging their psychometrically modified versions (see *Internal consistency* in the Supplement; *Supplementary Fig. **S4***), while the positive CT and SA results for PCL-R F1 (and less so for PCL-R F2) remained highly consistent when taking two alternative approaches to structural-data quality control (see *MRI* in the Supplement; *Supplementary Fig. **S5***).

**Figure 2.**
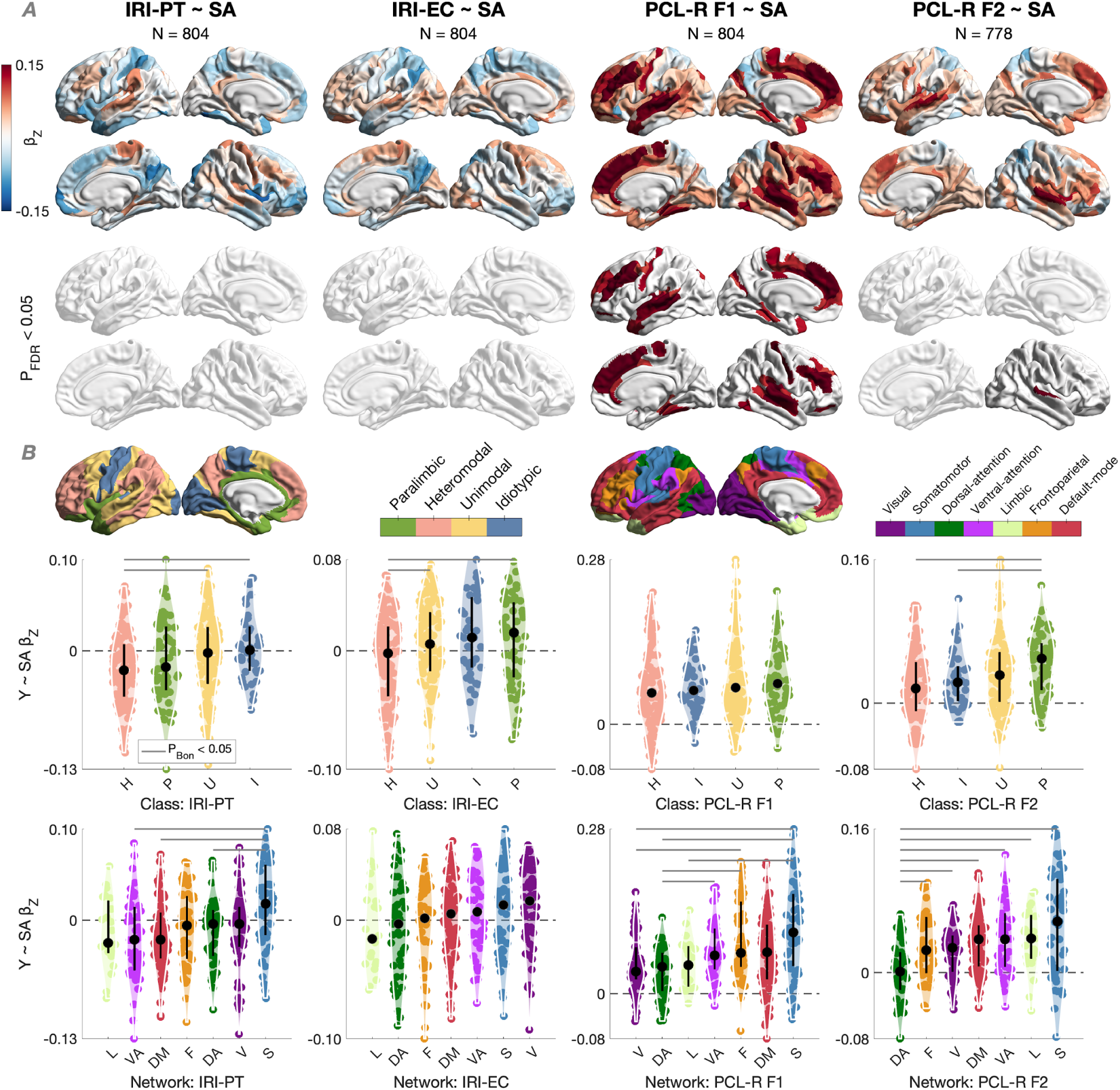
SA in relation to empathy and psychopathy (Q2). (A) Relationships of SA (positive, if any) with IRI-PT, IRI-EC, PCL-R F1, and PCL-R F2, controlling for age, IQ, and TIV in a robust linear regression with FDR correction. Grayed-out parcels are insignificant at P_FDR_ < 0.05. (B) Standardized betas across the cortex by Mesulam’s class and Yeo’s network, median-ordered and tested for distribution differences using Wilcoxon’s rank-sum test with Bonferroni’s correction within class (six comparisons) or network (21 comparisons).

We then tested for multivariate, predictive relationships of CT and SA with empathy and psychopathy using a train-test split and cross-validation (Q3). Most importantly, the univariate relationships of both CT and SA with PCL-R F1 (but not PCL-R F2) were corroborated: CT explained ∼6% of the out-of-sample variance in PCL-R F1, while SA explained ∼8% (*Fig. **3*** and *Supplementary Fig. **S6***).

**Figure 3.**
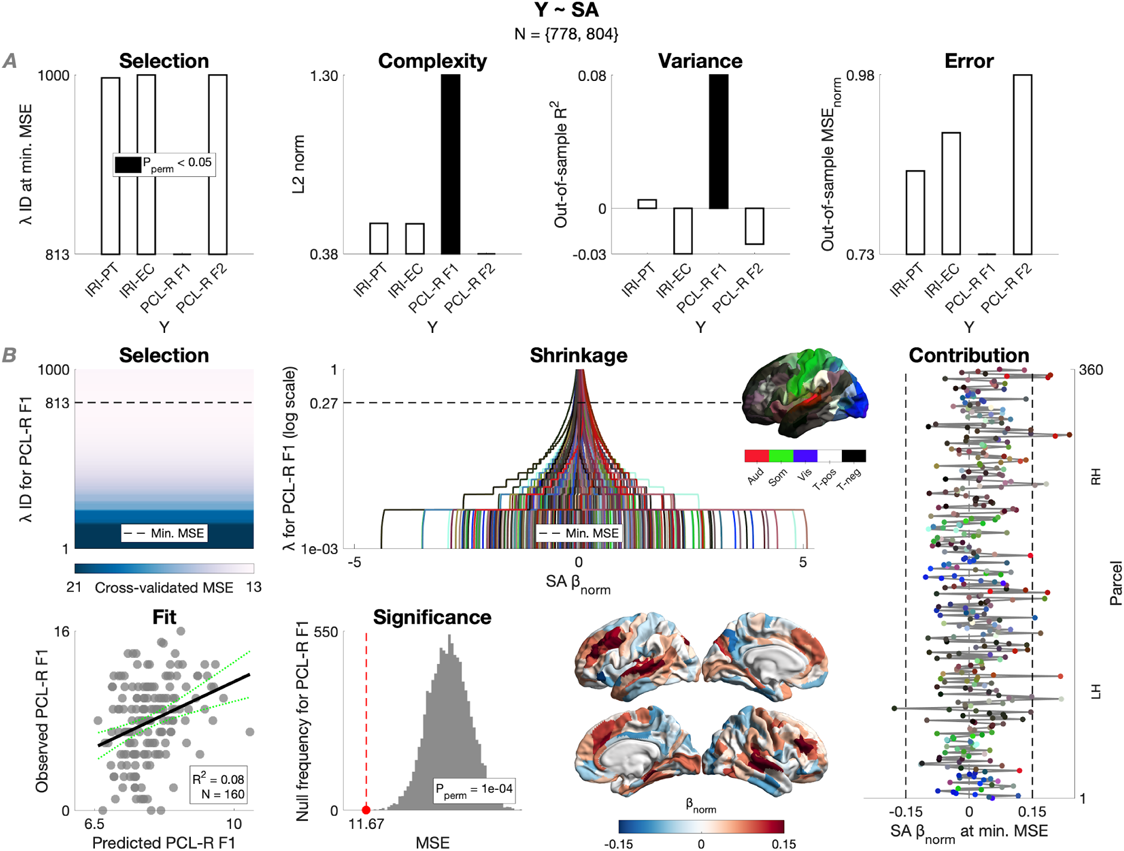
Multivariate prediction of empathy and psychopathy from SA (Q3). (A) For IRI-PT, IRI-EC, PCL-R F1, and PCL-R F2, we inform on: model selection using cross-validated ridge regression (i.e., lambda corresponding to the minimum cross-validated MSE at which the model was selected); model complexity (i.e., Euclidean norm of the final beta vector); variance explained (i.e., out-of-sample coefficient of determination); and prediction error (i.e., out-of-sample MSE divided by the maximum possible score and thus normalized). SA was corrected for age, IQ, and TIV separately in the training (N = 644/623) and test (N = 160/155) sets. Among the four variables, only PCL-R F1 was able to be predicted (R^2^ = 0.08 [95% CI: 0.02, 0.13], P_perm_ = 1e-04). (B) For PCL-R F1, we inform on model selection, beta shrinkage, final beta vector, predicted-observed fit, and significance based on permutation for out-of-sample MSE (N_perm_ = 10,000).

### Cortical structure by psychopathy group (Q4)

Next, we tested for global and regional differences in cortical structure by psychopathy group (Q4). There was no difference in CT (*Supplementary Fig. **S7***). In contrast, high-psychopathy men, compared to low-psychopathy men, had increased total SA. Regionally, there was an increase in 65 parcels (*Fig. **4A*** and *Supplementary Table **S10***); additionally controlling for race and substance use yielded highly similar results (*Supplementary Fig. **8***). More specifically, effect sizes across the cortex were largest in the paralimbic class and somatomotor network (positive median in both cases), with differentiation by both class (i.e., paralimbic > heteromodal) and network (e.g., somatomotor > dorsal-attention). We then computed spatial overlap between the FDR-corrected cluster of SA increases and meta-analytic brain-behavior data. First, using task-based activations underlying social-cognitive and social-affective processing (*Supplementary Fig. **S9***), the SA increases overlapped multiple times more with social-affective than social-cognitive clusters (*Fig. **4B***). Secondly, using task-based activations across 24 wide-ranging terms from Neurosynth (*Supplementary Fig. **S10***), the overlap was highest for affective/sensory terms (top two: “pain”, “auditory”) and lowest for visual terms (bottom two: “visual perception”, “visuospatial”) (*Fig. **4C***).

**Figure 4.**
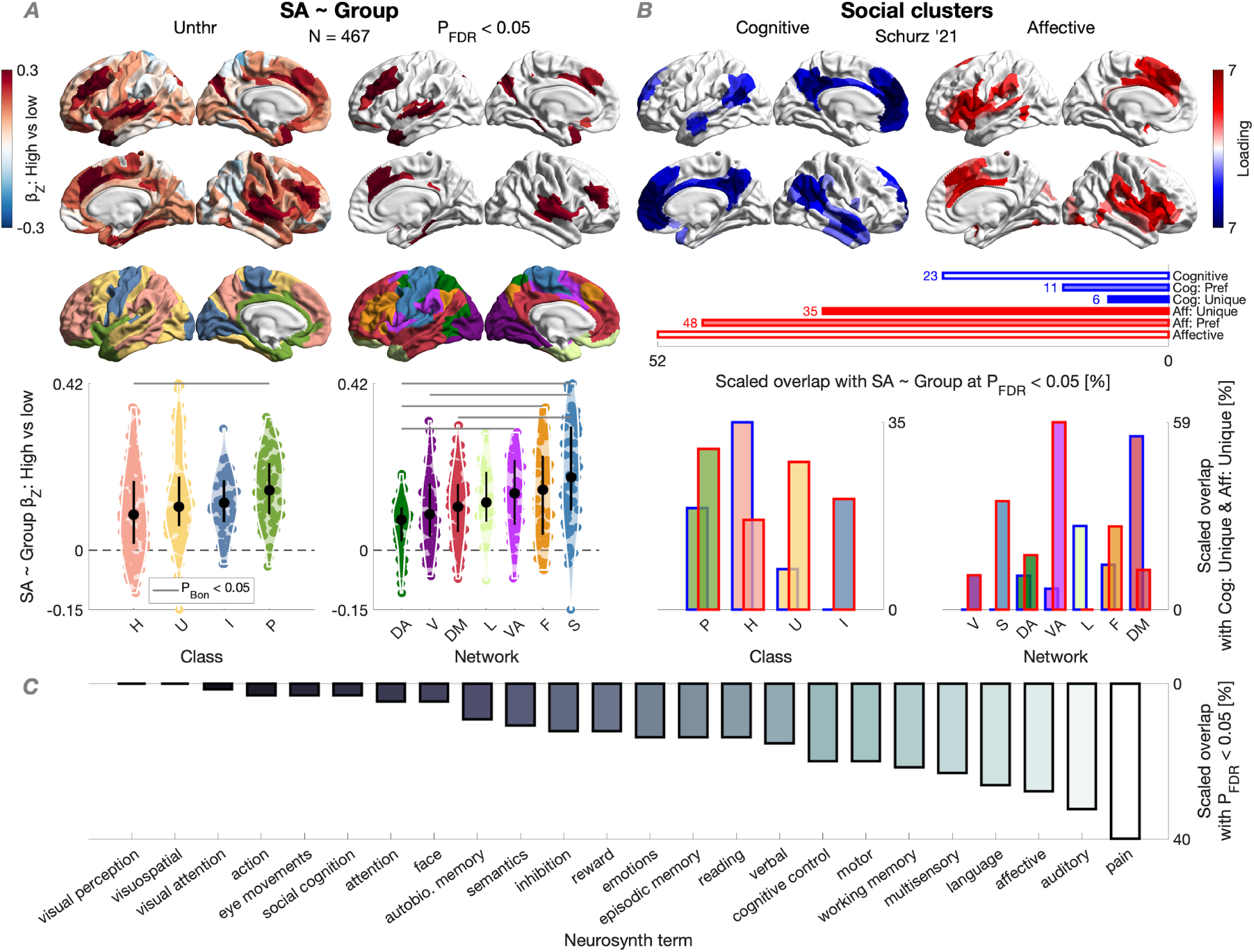
SA by psychopathy group (Q4). (A) Differences in SA by psychopathy group (high [N = 178] versus low [N = 289]), controlling for age, IQ, and TIV in a robust linear regression with FDR correction; 65 parcels showed an increase in high-psychopathy men. Across the cortex, standardized betas were median-ordered by class and network, and tested for distribution differences using Wilcoxon’s rank-sum test with Bonferroni’s correction within class (six comparisons) or network (21 comparisons). Total SA was increased in high-psychopathy men as well, controlling for the same covariates (β_Z_ = 0.25 [95% CI: 0.14, 0.36], P = 9e-06, adj. R^2^ = 0.67, Cohen’s D = 0.39). (B) Meta-analytic clusters of social-cognitive and social-affective processing across 130 studies (31). The FDR-corrected cluster of SA increases in high-psychopathy men overlapped multiple times more with social-affective than social-cognitive clusters across different social-cluster thresholds. “Cognitive” and “Affective” = baseline clusters; “Cog: Pref” and “Aff: Pref” = preferential clusters (including parcels with a higher loading compared to their baseline counterpart); “Cog: Unique” and “Aff: Unique” = unique clusters (including parcels not present in their baseline counterpart). Below, social-cluster overlap with the classes/networks (for the Cog: Unique and Aff: Unique clusters, which showed the highest proportional difference). (C) Overlap between the FDR-corrected cluster and Neurosynth clusters (102). For the overlap scaling, see the *Supplement*.

### Structural-covariance gradients by psychopathy group (Q5)

Finally, we tested structural-covariance gradients by psychopathy group (Q5). Taking the non-linear approach of diffusion-map embedding to decompose the high dimensionality of cortical structure in the total sample, we observed that the primary gradients of CT and SA traversed anterior-posterior axes, which were consistent with the HCP sample (*Fig. **5A-C*** and *Supplementary Fig. **S11***). We then tested for psychopathy-group differences in the gradients aligned to those in the total sample (*Fig. **5D***; for raw gradients, see *Supplementary Fig. **S12***). The primary gradient of CT was compressed in high-psychopathy men compared to low-psychopathy men, such that its distribution had a smaller range and was pulled toward the center. Such compression was not observed for SA (*Fig. **5B-C***). In sensitivity analyses for the CT gradient, compression was also observed for high versus moderate psychopathy (to a smaller extent); when lowering the high-psychopathy threshold; when matching the high- and low-psychopathy samples for size; and when using different alignment templates, including from the HCP sample (*Supplementary Fig. **S13***). Furthermore, aggregating gradient loadings by class and network revealed compression-oriented differences in the visual, limbic, and frontoparietal networks for both gradients, and further differences in the paralimbic class and dorsal-attention network for the CT gradient (*Fig. **6***).

**Figure 5.**
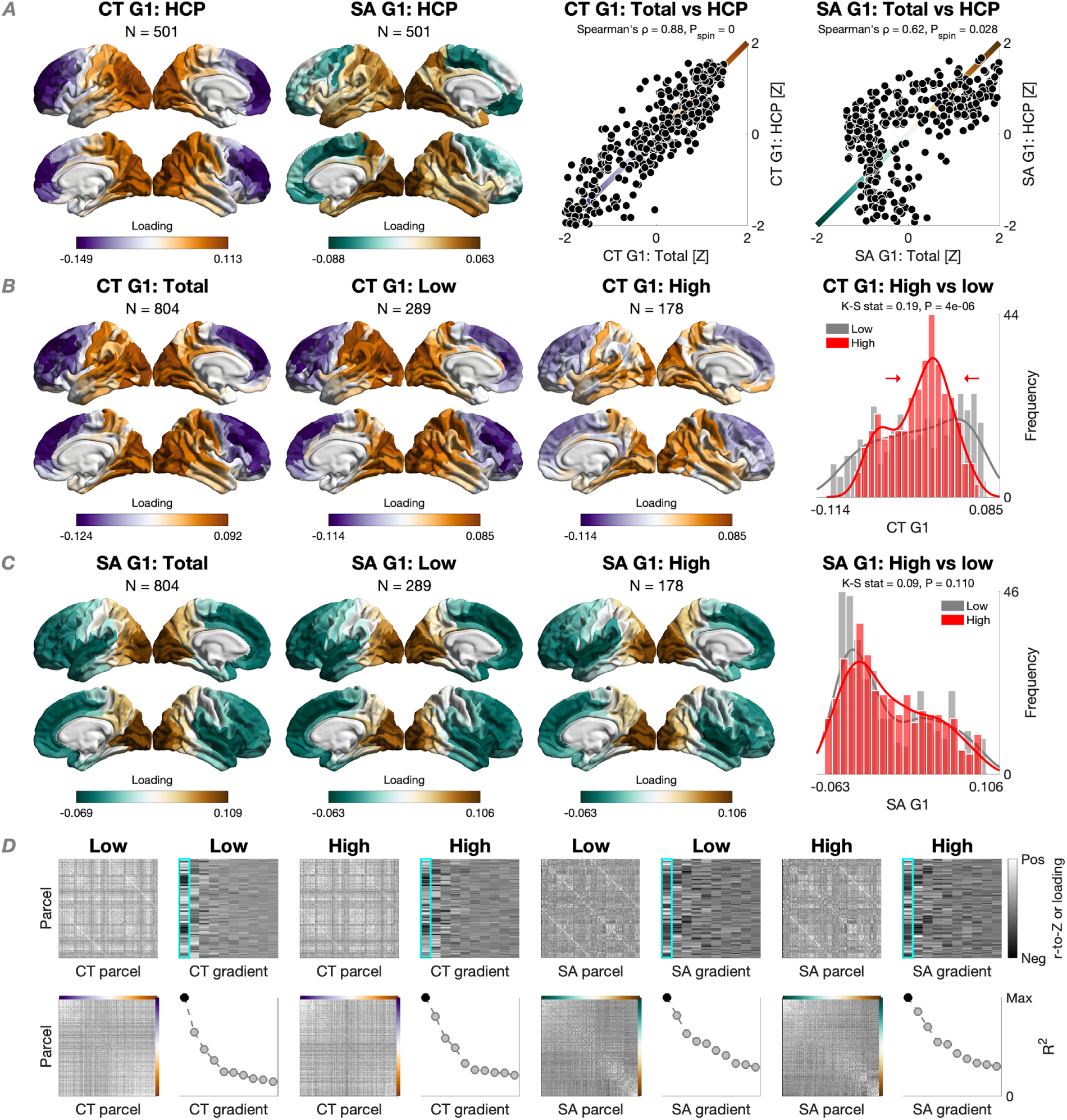
Macroscale organization of CT and SA by psychopathy group: Global analysis (Q5). (A) Primary gradients of CT and SA in the HCP sample (N = 501) and their spatial correlations with gradients in the total sample (N = 804) following spin permutation (N_perm_ = 1,000). In both datasets, CT was corrected for age and IQ while SA additionally for TIV. (B) Primary gradients of CT in the total, low-psychopathy, and high-psychopathy samples. The CT gradient was compressed in high-psychopathy men compared to low-psychopathy men using Kolmogorov-Smirnov’s test. (C) Primary gradients of SA in the three samples. The SA gradient did not differ by psychopathy group. (D) Consider the two-by-two left-hand tiles: Sample-specific covariance matrix (top left), array of the first 10 gradients (top right), covariance matrix ordered by the primary gradient (bottom left), and the first 10 gradients ordered by the proportion of the variance explained (i.e., scaled eigenvalues; bottom right). All matrices were set to the range [-0.5, 0.5]; all arrays were set to the minimum-maximum range.

**Figure 6.**
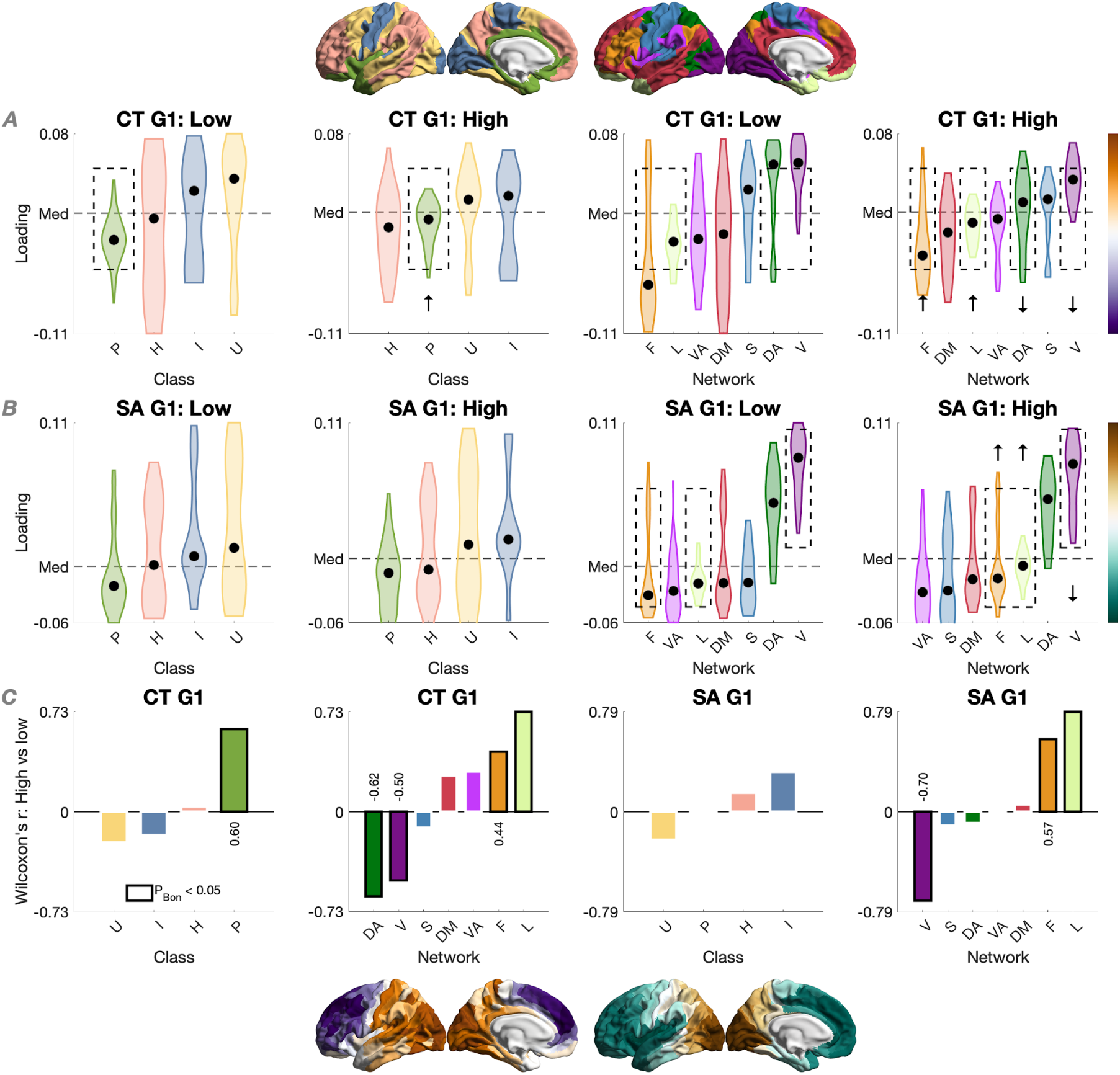
Macroscale organization of CT and SA by psychopathy group: Local analysis (Q5). (A) CT and (B) SA gradients by class and network in low-psychopathy (N = 289) and high-psychopathy (N = 178) men, median-ordered. Both gradients in low-psychopathy men, and the CT gradient in high-psychopathy men, traversed a frontoparietal-to-visual axis; the SA gradient in high-psychopathy men traversed a ventral-attention-to-visual axis instead. (C) Using Wilcoxon’s signed-rank test with Bonferroni’s correction within class (four tests) or network (seven tests), both gradients differed in high-psychopathy men compared to low-psychopathy men in a compression-oriented manner – where the class/network median was pulled toward the center (i.e., median across the classes/networks) – in the visual, limbic, and frontoparietal networks. The CT gradient further differed in this manner in the paralimbic class and dorsal-attention network.

## Discussion

A comprehensive, well-powered mapping of CT and SA in relation to empathy and psychopathy has been lacking. We addressed this gap in 804 incarcerated adult men through five overarching questions.

As expected, psychopathy had negative relationships with empathy (Q1). PCL-R F1 had a negative relationship with IRI-EC but not IRI-PT, while PCL-R F2 had a negative relationship with both IRI subscales. Controlling for the other subscale revealed statistically unique contributions of PCL-R F1 to IRI-EC and of PCL-R F2 to IRI-PT. In categorical analyses for high (i.e., clinical) levels of psychopathy, high-psychopathy men scored lower on both IRI subscales, but only the difference on IRI-EC (which was larger) proved unique. Evoking a “mirror-opposite” image with autism (7,105–109), this aligns with meta-analytic evidence that psychopathy is primarily associated with reduced affective empathy and empathic concern rather than cognitive empathy (10,11) (for a similar conclusion based on self-report data, see, e.g., (110)). We further add to this literature by revealing a pattern of unique relationships, suggesting that the psychopathic reduction in cognitive empathy – meta-analytically replicable also based on performance data (111–113) – may depend on the reduction in empathic concern, at least in part.

SA had positive relationships with psychopathy (Q2). This was observed for 103 out of 360 parcels for PCL-R F1 and three parcels for PCL-R F2. The superior-temporal/auditory cortex, playing a role in affective-speech processing (114), showed up for both PCL-R factors, with effect sizes across the cortex being largest in the paralimbic class and somatomotor network. These SA increases were in contrast to what we expected based on meta-analytic GMV reductions observed for male psychopathy (51), knowing that cortical GMV closely tracks SA genetically and phenotypically (115). It is important to note, however, substantial differences between our surface-based study and this meta-analysis: it included voxel-based-morphometry (VBM) studies across mixed (i.e., community and incarcerated) samples ranging from as few as N = 12 to 254 (total N = 519). In contrast to the PCL-R factors, we did not observe any relationship of SA with the IRI subscales, which raises questions about its reliability (116) and calls for measuring empathy beyond self-report in forensic neuroimaging. Regarding CT, we observed more circumscribed relationships with PCL-R F1 compared to SA (mostly positive) but not with any other behavioral variable tested. Together with our multivariate prediction of PCL-R F1 (but not PCL-R F2), where SA explained more out-of-sample variance than CT did (Q3), our results suggest that PCL-R F1 is more neuroanatomically distinctive than PCL-R F2, and that this is better captured by SA than CT. To stress the novelty, this is the first evidence of any relationship, positive or negative, between SA and psychopathy.

Corroborating the dimensional results, high-psychopathy men had increased SA (Q4). These increases spanned 65 parcels and also covered the superior-temporal/auditory cortex; additional parcels covered the insula, temporal pole, or dorsolateral, dorsomedial, and orbital prefrontal cortices. Again, effect sizes across the cortex were largest in the paralimbic class and somatomotor network. The paralimbic class has been hypothesized to be especially relevant to psychopathy under the “paralimbic-dysfunction” model (81). While the SA increases we observed indeed highlight paralimbic relevance, they do not readily align with the GMV reductions hypothesized by the model and commonly reported (17,51,80). What further highlights the relevance of (social-)affective/sensory regions to psychopathy in our study is the spatial overlap of the SA increases with meta-analytic task-based activations. In particular, the SA increases overlapped multiple times more with clusters of social-affective than social-cognitive processing across 130 studies (31), while across 24 wide-ranging brain-behavior meta-analyses (102), the overlap was highest for affective/sensory terms (“pain” and “auditory”). In contrast, we observed no psychopathy-group difference in CT. While these results agree with a higher sensitivity of SA than CT to broadly construed antisocial behavior, they again disagree with the effect direction. This is because reductions rather than increases in SA have been reported among antisocial individuals, although drawn exclusively (58) or largely (59) from the general rather than incarcerated population. It will thus be essential for future work to reconcile the SA increases we observed for psychopathy, dimensionally and categorically, with both the GMV reductions (meta-analytically) observed for psychopathy and the SA reductions observed for antisocial behavior (but see (50,80) for systematic reviews on psychopathy noting GMV increases). To enhance the developmental framing of the SA increases observed here, considered should be the cellular and physical mechanisms driving cortical expansion, including neural-progenitor proliferation, tangential neuronal migration, and mechanical stress (117–119).

Finally, high-psychopathy men had a compressed macroscale organization of CT (Q5). The primary gradient in the total sample traversed an anterior-posterior axis for CT (as reported across the sexes in the HCP, likely mirroring the temporal sequence of neurogenesis (67)) and a similar axis for SA (as reported in the genomic literature (120)). Both axes were consistent with the male HCP sample, more evidently the CT axis, suggesting its normativity. When testing for psychopathy-group differences along these axes, we observed a globally compressed gradient of CT but not SA. At the class and/or network level, high-psychopathy men further showed compression-oriented differences for both gradients, with converging and largest differences in the limbic network. CT gradients are known to differ across (71–73) and within (79) major psychiatric conditions, as is the primary functional gradient in terms of compression (74–76). We provide the first evidence that high-psychopathy individuals may exhibit similar macroscale properties of the cortex, with reduced differentiation between anterior/transmodal regions (e.g., frontoparietal) and posterior/unimodal regions (e.g., visual) – which anchored the ends of our CT gradient. It is an open question whether such reduced differentiation reflects disrupted integration and segregation from a connectomic perspective (78,121,122), which sets the stage for probing psychopathy-related differences along the unimodal-transmodal axis itself (70). This axis not only recapitulates the anterior-posterior axis of CT (67) but may be clinically compressed along with the CT axis, as suggested for schizophrenia (79).

There are limitations to our study. For instance, a performance test of empathy could have yielded additional insights. Indeed, our use of the IRI cannot be conclusive given the questionable correspondence between self-reported and tested empathy (123), and the potential for social-desirability bias and metacognitive deficits in the incarcerated population. As above, our results may not generalize to community (124) but also to female samples, with there being sex/gender differences in empathy (125–127) and cortical structure (128), beyond psychopathy. Recruiting from the general and incarcerated female populations should thus be prioritized. Furthermore, the roles of other macrostructural (e.g., folding and curvature) as well as microstructural (e.g., diffusion-based) and subcortical indices remain to be elucidated.

In conclusion, high-psychopathy men had reduced empathic concern, increased SA, and a compressed macroscale organization of CT, suggesting specific co-occurring alterations in empathy and cortical structure. Future work should build upon these novel insights in both the general and incarcerated populations to inform the treatment of psychopathic traits such as reduced empathy.

## Supporting information

Radecki et al. (2025) - Supplement

Radecki et al. (2025) - Supplement (Excel)

## Acknowledgments and Disclosures

Author contributions are as follows: Conceptualization: M.A.R., E.S., G.L., G.H., S.P., P.P., K.A.K., & L.C. Data processing: M.A.R., J.M.M., K.A.H., D.D.S., & L.C. Data analysis: M.A.R. & L.C. Data visualization: M.A.R. Writing – original draft: M.A.R. Writing – review and editing: All authors. Funding and data acquisition: C.L.H., S.P., J.D., & K.A.K. Supervision: P.P., K.A.K., & L.C.

Some of the generated data and code will be shared online upon publication (https://github.com/MARadecki/EmpathyPsychopathy/). Regarding data from the incarcerated sample, please contact K.A.K. Human Connectome Project data are available online (https://www.humanconnectome.org/). The following resources are further available online: meta-analytic brain-behavior data from Schurz et al. (https://osf.io/pav27/ (31)) and Neurosynth (https://neurosynth.org/ (102)); volume-to-surface mapping, parcellation, and/or gradient mapping as part of neuromaps (https://neuromaps-main.readthedocs.io/ (129)), ENIGMA Toolbox (https://enigma-toolbox.readthedocs.io/ (130)), and BrainSpace (https://brainspace.readthedocs.io/ (131)).

M.A.R. is supported by the Autism Research Trust Fund. All research at the Department of Psychiatry in the University of Cambridge is supported by the NIHR Cambridge Biomedical Research Centre (NIHR203312) and the NIHR Applied Research Collaboration East of England. The views expressed are those of the author(s) and not necessarily those of the NIHR or the Department of Health and Social Care.

This study was funded by the National Institute of Mental Health through grant numbers R01 MH070539 (PI: Kiehl), R01 MH114028 (PI: Harenski), and R01 MH071896 (PI: Kiehl), the National Institute on Drug Abuse (NIDA) through grant numbers R01 DA026505 (PI: Kiehl), R01 DA026964 (PI: Kiehl), and R01 DA020870 (PI: Kiehl), the National Institute of Child Health and Human Development (NICHD) through grant number R01 HD092331 (PI: Kiehl), the National Institute of Neurological Disorders and Stroke (NINDS) through grant number R01 NS126742 (PI: Kiehl), and the Italian Ministry of Education and Research through grant number PRIN2020 2020WSCSLZ (“Hot for genes – the role of brain gene expression in identifying antisocial developmental trajectories and malleable risk factors for preventive interventions”; PI: Pellegrini).

Data were provided [in part] by the Human Connectome Project, WU-Minn Consortium (Principal Investigators: David Van Essen and Kamil Ugurbil; 1U54MH091657) funded by the 16 NIH Institutes and Centers that support the NIH Blueprint for Neuroscience Research; and by the McDonnell Center for Systems Neuroscience at Washington University.

We thank Matthias Schurz for making meta-analytic data from (31) publicly available. All authors report no biomedical financial interests or potential conflicts of interest.

