## Supplementary material for "Cortical structure in relation to empathy and psychopathy in 800 incarcerated men": Radecki et al. (2025) - Supplement

Supplement including

Figs. S1-S13 and Tables S1-S5

### Supplementary methods

#### MRI

Each of the 912 T1-weighted MRI scans underwent the standard recon-all pipeline in FreeSurfer version 7.4.1 (https://surfer.nmr.mgh.harvard.edu/ (1)). To delineate 360 regions, the output was parcellated by resampling the HCP-MMP1.0 template (2) in the fsaverage space (3) to the native space via FreeSurfer's surface-based registration. For quality control, we applied thresholding by the Euler number (4), defined as the total number of topological defects in the cortical surface prior to fixing in the recon-all pipeline (5). In particular, we excluded participants whose Euler number was greater than 3 median absolute deviations (MADs) above the median (N = 105 out of 912; ~12%). Since the Euler number has no universally accepted threshold – nor is there a "gold standard" for the quality control of FreeSurfer output in general (6) – we ran two sensitivity analyses for some of the main structural results in the total sample (i.e., results at $P_{FDR} < 0.05$ (7)). First, we used a much more conservative threshold (> 2 MADs above the median, excluding N = 154, i.e., ~17%; note that this threshold in Bethlehem et al. (5) excluded ~9-10% of scans per dataset, while we already surpassed this percentage at > 3 MADs). Secondly, we controlled for the Euler number as a covariate instead of applying thresholding based on it. Both sensitivity analyses were supplemented with tests of spatial correspondence with the main whole-cortical maps of effect size (standardized beta) using Spearman's correlation and spin permutation (8,9), as implemented in the ENIGMA Toolbox version 2.0.3 (https://enigma-toolbox.readthedocs.io/ (10)) under MATLAB version R2020b (The MathWorks Inc., Natick, MA; https://www.mathworks.com/).

Structural-MRI data in the Human Connectome Project (HCP) were acquired and underwent the "minimal-preprocessing" pipeline as described in (11). FreeSurfer output was parcellated in FreeSurfer version 6.0.0 using the same HCP-MMP1.0 template and code as above, while total intracranial volume (TIV) and the Euler number were extracted from aseg.stats files.

All subsequent plotting of the cortical data was done using the ENIGMA Toolbox and the HCP-MMP1.0 template in a 32k fs_LR space (https://balsa.wustl.edu/file/show/3VLx/).

#### Internal consistency

Internal consistency of the Interpersonal Reactivity Index (IRI) and Psychopathy Checklist-Revised (PCL-R; using all available data) was analysed using the omega function in the psych package version 2.5.3 in R version 4.5.0 (12). Internal consistency was acceptable for both the Perspective Taking (IRI-PT; unidimensional McDonald's $\omega_t$ = 0.766) and Empathic Concern (IRI-EC; unidimensional McDonald's $\omega_t$ = 0.799) subscales. In addition, aiming to improve their reliability for sensitivity analyses with neuroimaging data (13), we calculated IRI-PT and IRI-EC scores using positively scored items only, since the reverse ones tend to show smaller factor loadings (see (14) for a meta-analysis; see (15) for an example study in a similarly sized sample of incarcerated males that showed acceptable internal consistency for IRI-PT and IRI-EC across all items, and acceptable-to-good internal consistency across positively scored items). Here, internal consistency improved to good for IRI-PT (unidimensional McDonald's $\omega_t$ = 0.802), while it remained acceptable for IRI-EC (unidimensional McDonald's $\omega_t$ = 0.759).

Internal consistency was good for the PCL-R (unidimensional McDonald's $\omega_t$ = 0.816; bidimensional McDonald's $\omega_t$ = 0.836) and acceptable for both the PCL-R F1 (unidimensional McDonald's $\omega_t$ = 0.772) and PCL-R F2 (unidimensional McDonald's $\omega_t$ = 0.703) items.

**Covariates**

Crime type was self-reported and classified similarly to (16). In particular, we classified the following crimes as violent: assault/battery, assaulting a police officer, attempted murder, child abuse/neglect, child sexual assault, domestic assault/battery, manslaughter, murder, rape/sexual assault, and robbery. We classified the following crimes as non-violent: arson, burglary, drug distribution, drug possession, escape, failure to appear in court, fraud/forgery, parole/probation violation, pimping, prostitution, resisting arrest, theft (of any monetary value), and vandalism.

Full-scale intelligence (i.e., IQ) was estimated with the Vocabulary and Matrix Reasoning subtests of the Wechsler Adult Intelligence Scale, third edition (17) (N = 705) or the same subtests of the Wechsler Abbreviated Scale of Intelligence, second edition (18) (N = 99). The Vocabulary score ranges 0-66 points and the Matrix Reasoning score ranges 0-26 points, with higher scores indicating higher intelligence. We summed the two scaled scores and converted them into an IQ estimate.

Race was self-reported as "American Indian or Alaskan Native", "Asian", "Black or African American", "Native Hawaiian or other Pacific Islander", "White", or "More than one race". Given that ~68% of the total sample self-identified as White, race was collapsed into White versus non-White. This variable was available for N = 789 in the total sample (N = 804).

Substance use was measured with the Addiction Severity Index, fifth edition (19). In particular, we identified the total number of years of regular substance use (with "regular" meaning three or more times per week for a minimum of three months) and collapsed this number across substances. We also report this variable that was divided by age (to correct for opportunity to use) and then square-root-transformed (to correct for skewness). Both variables were available for N = 748.

See *Table* ***1*** for IQ, race, and substance use by sample (total, low-psychopathy, and high-psychopathy).

**Mesulam's classes and Yeo's networks**

We included four laminar-differentiation classes according to Mesulam (20). These distinguish paralimbic areas (e.g., cingulate cortex), heteromodal (higher-order) association areas (e.g., medial prefrontal cortex), unimodal (modality-specific) association areas (e.g., lateral occipital cortex), and idiotypic (primary) areas (e.g., visual cortex). For class assignments in the HCP-MMP1.0 atlas, we followed Dorfschmidt et al. (21,22). Four parcels without an assignment (L_Pir, L_H, R_Pir, and R_H, for the piriform cortex and hippocampus) were treated as missing.

We also included seven intrinsic-connectivity networks according to Yeo et al. (23): visual, somatomotor, dorsal-attention, ventral-attention, limbic, frontoparietal, and default-mode. For network assignments in the HCP-MMP1.0 atlas, we parcellated the fsaverage5 template (https://github.com/ThomasYeoLab/CBIG/) based on the mode (as in prior work (24)). Four parcels without an assignment (the same ones as above) were treated as missing.

**Multivariate prediction**

To predict empathy and psychopathy from CT and SA in a machine-learning context (25), we used ridge regression as implemented in fitrlinear function under MATLAB R2020b. First, training and test sets were randomly partitioned with an 80-20 split under the default seed together for IRI-PT, IRI-EC, PCL-R total, and PCL-R F1 (N = 644/160), and then for PCL-R F2 (N = 623/155), given that the latter had missing observations. CT was then corrected for age and IQ, while SA was additionally corrected for TIV in a robust linear regression, from which raw residuals were extracted. This was done separately in the training and test sets to avoid data leakage (26). Predictor data (i.e., CT and SA) were then normalized to the range [0, 1] using the minimum and maximum values in the training set to ensure consistent scaling across the training and test sets. Ridge regularization (i.e., L2 penalty) was then initiated using 10-fold cross-validation with 1,000 logarithmically spaced lambdas, ranging [0.001, 1], with the least-squares learner and the LBFGS solver under the default seed. Across the lambdas, we selected the one corresponding to the minimum cross-validated mean squared error (MSE) to balance model complexity and generalization. Next, we computed predicted scores, L2 norm (as the Euclidean norm of the final beta vector), sum of squared errors (SSE), total sum of squares (SST), out-of-sample coefficient of determination ($R^2$), and out-of-sample MSE, as follows:

$$\widehat{Y} = \text{Intercept}_{\text{training}} + (X_{\text{test}} \times \beta_{\text{training}})$$

$$\text{SSE} = \sum_{i=1}^{N} (Y_{\text{test,i}} - \widehat{Y}_{\text{test,i}})^2$$

$$\text{SST} = \sum_{i=1}^{N} (Y_{\text{test,i}} - \overline{Y}_{\text{test}})^2$$

$$R^2 = 1 - \frac{SSE}{SST}$$

$$\text{MSE} = \frac{1}{N} \sum_{i=1}^{N} (Y_{\text{test,i}} - \widehat{Y}_{\text{test,i}})^2$$

where $N$ denotes the number of participants, $i$ the participant, Y their observed score, $\widehat{Y}$ their predicted score, and $\overline{Y}$ mean observed score. For the model yielding a positive $R^2$, we inferred significance via permutation for the MSE by shuffling CT or SA labels ($N_{\text{perm}}$ = 10,000). Finally, we computed the 95% confidence interval for the $R^2$ in the significant model by bootstrapping the test data ($N_{\text{boot}}$ = 10,000), recomputing the $R^2$ for each sample, and using the percentile method for the interval boundaries.

**Meta-analytic task-based activations**

To enhance the interpretability of structural differences by psychopathy group in a framework distinguishing social-cognitive and social-affective processing, we used the volumetric meta-analysis by Schurz et al. (27,28). This meta-analysis leveraged task-based fMRI and PET activation across 188 studies with a total N = 4,207, applying a voxel-wise threshold of $P < 0.005$ and a cluster-extent threshold of 10 voxels, which were found to be optimal in balancing sensitivity and specificity, and approximately corresponded to a corrected threshold of $P < 0.05$ in the original studies. Among the three hierarchically derived brain-activation clusters (cognitive, affective, and intermediate), we used the cognitive and affective ones ("cluster 1" and "cluster 3"), based on 57 and 73 studies, respectively. The cognitive cluster represented "predominantly cognitive processes, which are engaged when mentalizing requires self-generated cognition decoupled from the physical world", while the affective cluster represented "more affective processes, which are engaged when we witness emotions in others based on shared emotional, motor, and somatosensory representations" (27, p. 294).

To further explore the broader psychological relevance of the psychopathy-group differences, we used 24 meta-analytic clusters from Neurosynth version 0.7, an automated meta-analytic referencing of task-based fMRI activation (https://neurosynth.org/ (29)). In particular, we used "association tests", which represent Z-scores from a two-way ANOVA that tests for the presence of a non-zero association between term use and voxel activation, with the results being corrected for multiple testing at $P_{FDR} < 0.01$. These 24 terms were selected for breadth and consistency with prior work (e.g., (30-32)).

Both the Schurz et al. (henceforth, "social") and Neurosynth clusters in MNI space were projected onto a 32k fs_LR surface via registration fusion (33,34) and parcellated in the HCP-MMP1.0 atlas using neuromaps version 0.0.5 (https://netneurolab.github.io/neuromaps/ (35)) in Python version 3.11.9 (https://www.python.org/). Since our characterization boiled down to computing spatial overlap with the social and Neurosynth clusters, and some of these clusters covered more than 50% of all parcels, each meta-analytic cluster was thresholded by the mean across all parcels for increased specificity (i.e., all loadings below the mean were nullified). For the social clusters, this yielded the "baseline", partly overlapping clusters ("Cognitive" and "Affective"). To better disentangle their preferential and unique profiles, respectively, they were thresholded by subtracting one from another and including the positive output ("Cog: Pref" and "Aff: Pref") as well as by identifying parcels that were not included in the baseline counterpart ("Cog: Unique" and "Aff: Unique"). The social clusters at each derivation step are shown in *Supplementary Fig.* ***9***, while the Neurosynth clusters are shown in *Supplementary Fig.* ***10***.

We computed spatial overlap at three levels of characterization. First, we computed spatial overlap between the psychopathy-group differences and social clusters, where scaling was done by dividing the number of overlapping parcels by the number of FDR-corrected parcels (for the psychopathy-group comparison) to indicate what proportion of the latter fell into each social cluster. Secondly, for the social clusters that yielded the highest proportional difference, we computed spatial overlap with Mesulam's classes and Yeo's networks, where scaling was done by dividing the number of overlapping parcels by the number of class/network parcels. Finally, we computed the spatial overlap between the psychopathy-group differences and Neurosynth clusters, where scaling was done as for the social clusters.

**Structural-covariance gradients**

To investigate macroscale structural organization as structural-covariance gradients, first, in the total sample (N = 804) and the HCP sample (N = 501), CT was corrected for age and IQ, while SA was additionally corrected for TIV in a robust linear regression, from which raw residuals were extracted. Sample-specific structural-covariance matrices were then computed using Pearson's r with Fisher transformation. To derive gradients from these matrices, we used the standard set of parameters in BrainSpace version 0.1.10 (https://brainspace.readthedocs.io/ (36)) under MATLAB version R2020b, in line with the extensive multimodal structural literature (e.g., (31,32,37-40)): diffusion-map embedding as the non-linear dimensionality-reduction technique (41), normalized angle as the kernel, 90% as the matrix sparsity, and the two hyperparameters, alpha and diffusion time, set at 0.5 and 0, respectively. Using CT and SA corrected in the total sample, gradients in low-psychopathy (N = 289) and high-psychopathy (N = 178) men were then derived in the same way. To ensure that they traversed the same axes and were thus directly comparable, we aligned them using the Procrustes method to those in the total sample using 10 gradients as the reference. In all downstream analyses, we included only the primary gradients, as they capture the highest proportion of variance akin to the linear principal-components analysis. The primary gradient of CT explained the following proportions of variance: ~22% in the total and low-psychopathy samples, ~18% in the high-psychopathy sample, and ~24% in the HCP sample. The primary gradient of SA explained ~14% of the variance in each of the incarcerated samples and ~13% in the HCP sample.

While the above gradients were included in the main analysis, we derived additional ones for sensitivity analyses by psychopathy group. First, we compared high- and low-psychopathy men with moderate-psychopathy men (i.e., all remaining participants, scoring PCL-R > 20 and PCL-R < 30; N = 337); gradients in this "moderate" sample were also aligned to those in the total sample. Secondly, we compared high- and low-psychopathy men by including more participants in the (smaller) high-psychopathy group, starting at PCL-R ≥ 27, "(…) which is 1 SEM unit below the conventional PCL-R threshold for psychopathy of 30" ((42), p. 375). Thirdly, we compared the high- and low-psychopathy samples after matching them for size by randomly subsampling the (larger) low-psychopathy sample or by bootstrapping the (smaller) high-psychopathy sample. In both cases, this was done 1,000 times to derive gradients for each sample and to compute the median gradient for the final comparison. Lastly, we compared high- and low-psychopathy men using different templates for Procrustes alignment, given the ongoing discussion on case-control comparisons in gradient space (43). In particular, first, we aligned the (raw) gradients in high-psychopathy men to the (raw) gradients in low-psychopathy men; secondly, we aligned gradients in both groups to those in the HCP sample.

**Statistical analysis**

Statistical analysis was conducted in MATLAB version R2020b and organized according to the five overarching questions: purely behavioral (dimensional and categorical) analyses for Q1; univariate (associative) brain-behavior analyses for Q2; multivariate (predictive) brain-behavior analyses for Q3; categorical analyses of cortical structure for Q4; and categorical analyses of structural-covariance gradients for Q5. All analyses apart from the predictive ones for Q3 were conducted using two-tailed tests to evaluate the null hypothesis of no effect against the alternative of any effect, positive or negative. At the request of a reviewer to state hypotheses in addition to the questions, we articulated the effect direction we had expected for some of the questions based on the reviewed meta-analytic literature: negative relationships of psychopathy with empathy (at least IRI-EC, e.g., IRI-EC by psychopathy group (44,45); Q1) and with cortical structure (at least SA, e.g., SA by psychopathy group (46); Q2 and Q4). Negative SA-psychopathy relationships were expected based on the unanimous meta-analytic reductions in cortical gray-matter volume (GMV) observed for male psychopathy (46), knowing that cortical GMV closely tracks SA but not CT, with phenotypic correlations up to ~0.9 for SA but only ~0.3 for CT (47).

Regarding covariates, all analyses (including CT analyses) controlled for (or were corrected for) age and IQ, while all SA analyses additionally controlled for (or were corrected for) TIV, in a robust linear regression. Regarding effect sizes, for all analyses, we reported standardized betas and/or Cohen's Ds. We recommend the reader to interpret these in line with the standard guidelines (48): 0.1, 0.3, and 0.5 (standardized beta) and 0.2, 0.5, and 0.8 (Cohen's D) for a small, medium, and large effect, respectively.

To address Q1, we first tested for relationships of psychopathy (PCL-R total, PCL-R F1, and PCL-R F2, as independent variables) with empathy (IRI-PT and IRI-EC, as dependent variables), controlling for age and IQ in a robust linear regression in the total sample (N = 804, apart from PCL-R F2; N = 778). We then tested IRI-PT and IRI-EC by psychopathy group (high versus low psychopathy; N = 178 versus 289, respectively), controlling for the same covariates. To further investigate statistically unique contributions of psychopathy to IRI-PT versus IRI-EC, we repeated both the dimensional and categorical analyses while additionally controlling for the other IRI subscale. In further sensitivity analyses by psychopathy group, we additionally controlled for race and total years of substance use. All the analyses were corrected for multiple testing across the IRI subscales using Bonferroni's correction.

To address Q2, we tested for relationships of cortical structure (CT and SA, as independent variables) with empathy and psychopathy (IRI-PT, IRI-EC, PCL-R total, PCL-R F1, and PCL-R F2, as dependent variables), controlling for age and IQ (with CT), and additionally for TIV (with SA), in a robust linear regression in the total sample. All these analyses were corrected for multiple testing across the 360 parcels using FDR correction. Standardized betas across the cortex were further aggregated by Mesulam's class and Yeo's network, median-ordered, and tested for distribution differences using Wilcoxon's rank-sum test with Bonferroni's correction within class (six comparisons) or network (21 comparisons).

To address Q3, which complemented the univariate analyses for Q2, we leveraged a multivariate framework to predict empathy and psychopathy from cortical structure (corrected for the same covariates) using ridge regression in the total sample. Details on this framework are provided in *Multivariate prediction* above.

To address Q4, we tested for global and regional differences in cortical structure (CT and SA, as dependent variables) by psychopathy group, controlling for age and IQ (with CT), and additionally for TIV (with SA), in a robust linear regression. For the global analyses, both mean CT and total SA were defined at the vertex level. All the regional analyses were corrected for multiple testing across the 360 parcels using FDR correction. In sensitivity analyses for those that yielded significant differences (at $P_{FDR} < 0.05$), we additionally controlled for race and total years of substance use. Whole-cortical standardized betas in the regional analyses were further aggregated by class/network, median-ordered, and tested for distribution differences using Wilcoxon's rank-sum test with Bonferroni's correction within class (six comparisons) or network (21 comparisons).

Finally, to address Q5, we tested for differences in CT and SA gradients by psychopathy group. First, gradient consistency between the total and HCP samples was evaluated via spatially correlation with Spearman's correlation and spin permutation. Gradients in high- and low-psychopathy men, aligned to those in the total sample, were then compared at a global level using a two-sample Kolmogorov-Smirnov's test to non-parametrically infer if they were drawn from the same distribution. Further, these gradients were compared at the class and network level using Wilcoxon's signed-rank test for significance and Wilcoxon's r for effect size (Wilcoxon's $r = Z / \sqrt{N}$), with Bonferroni's correction within class (four tests) or network (seven tests).

### Supplementary results

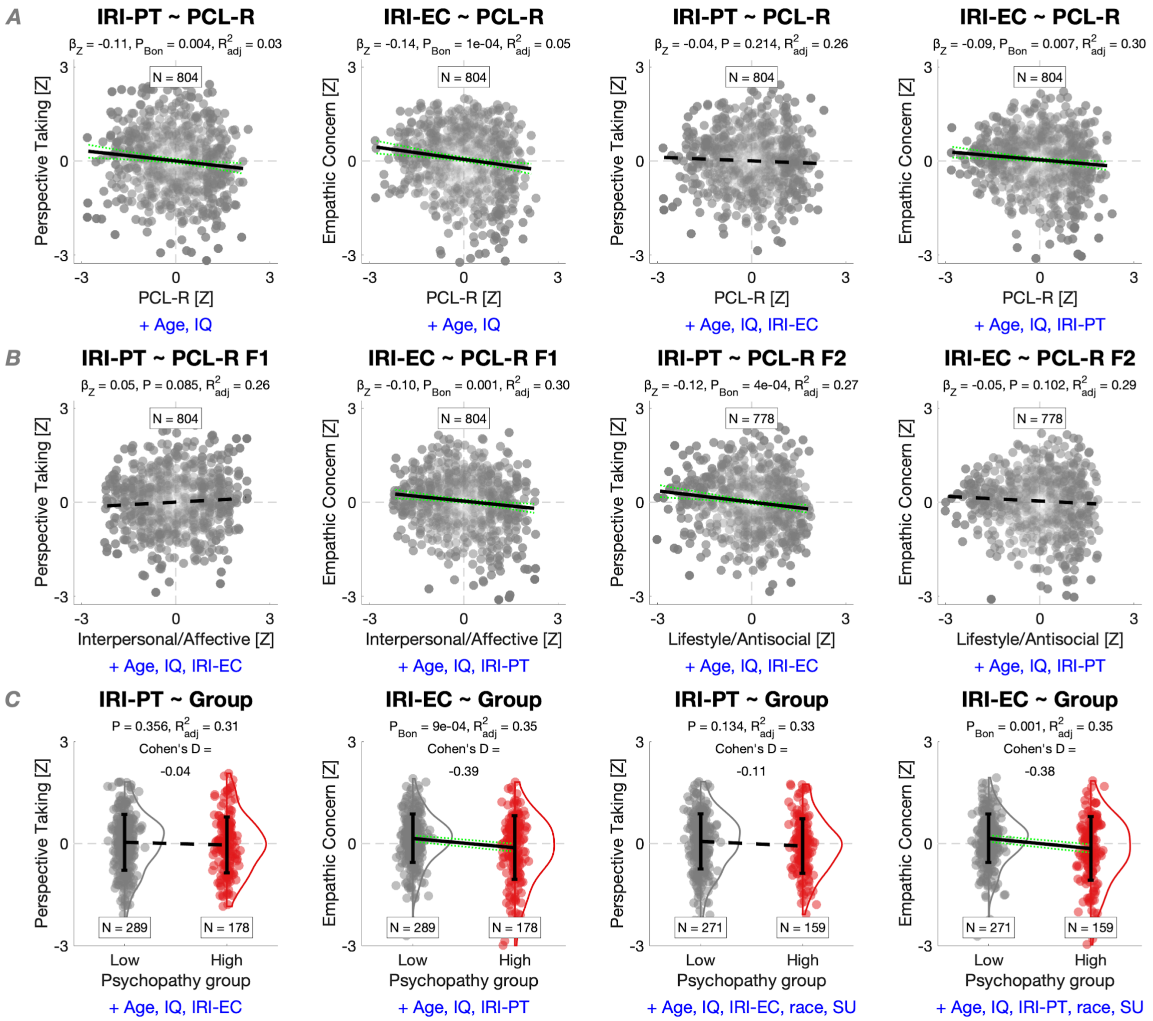

**Figure S1. Psychopathy in relation to empathy: Sensitivity analyses.** (A) PCL-R total in relation to IRI-PT and IRI-EC, controlling for age and IQ in a robust linear regression. In addition, models 3 and 4 controlled for the other IRI subscale. (B) PCL-R F1 and PCL-R F2 in relation to IRI-PT and IRI-EC, controlling for age, IQ, and the other IRI subscale. (C) IRI-PT and IRI-EC by psychopathy group, controlling for age, IQ, and the other IRI subscale. In addition, models 3 and 4 controlled for race and total years of substance use – two variables on which the groups differed. Across the panels, Bonferroni's correction was applied across the IRI subscales.

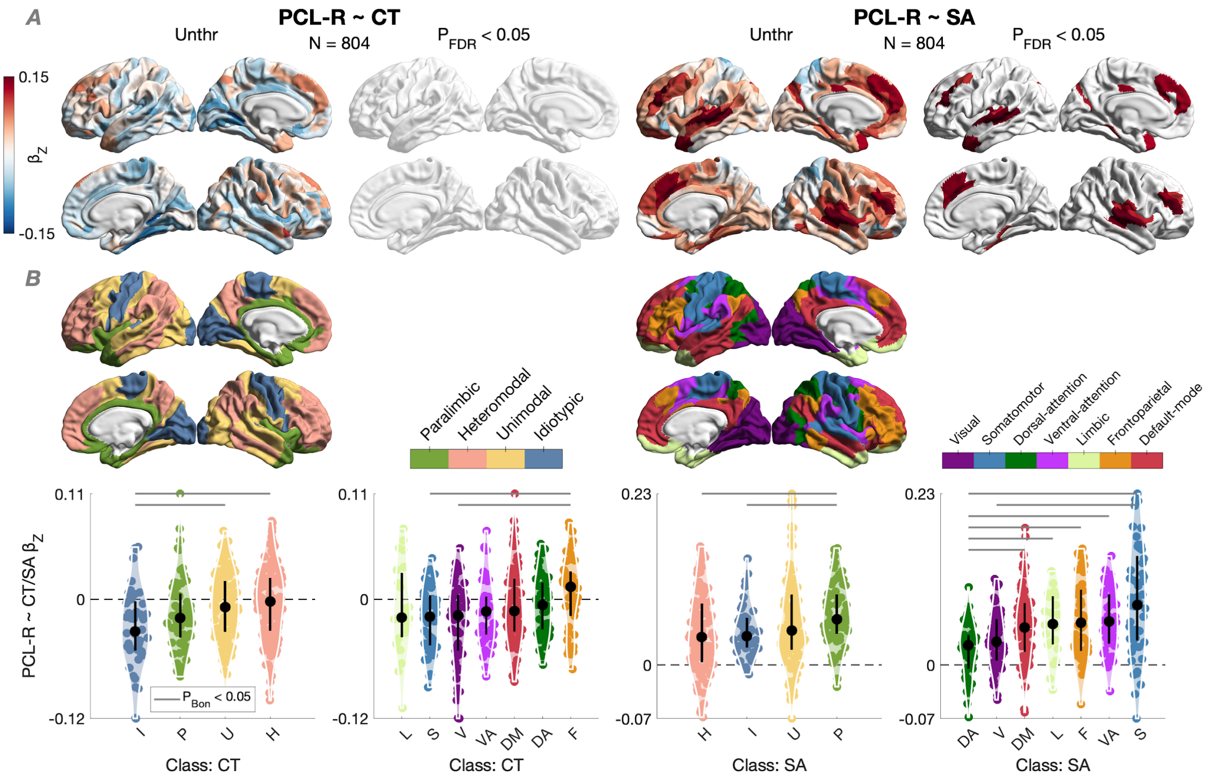


**Figure S2. CT and SA in relation to psychopathy (PCL-R total).** (A) Relationships of CT and SA with PCL-R total, controlling for age and IQ in a robust linear regression with FDR correction. In addition, the SA model controlled for TIV. N = 51 parcels had a positive relationship with PCL-R total, which is fewer than when testing SA by psychopathy group (i.e., N = 65; *Fig. **4***). (B) Standardized betas across the cortex by Mesulam's class and Yeo's network, median-ordered and tested for distribution differences using Wilcoxon's rank-sum test with Bonferroni's correction within class (six comparisons) or network (21 comparisons).

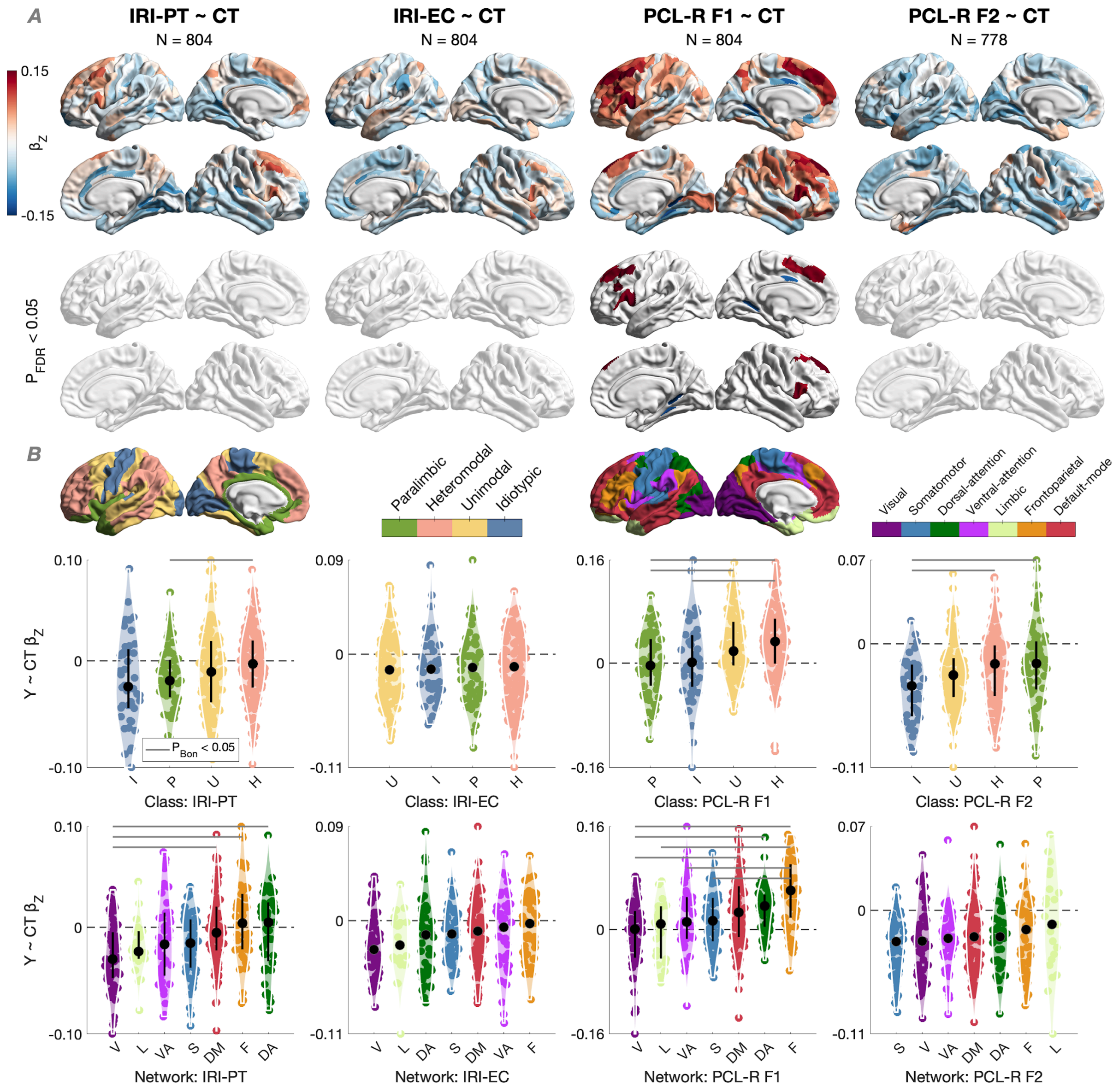


**Figure S3. CT in relation to empathy and psychopathy (PCL-R factors).** (A) Relationships of CT (mostly positive, if any) with IRI-PT, IRI-EC, PCL-R F1, and PCL-R F2, controlling for age and IQ in a robust linear regression with FDR correction. (B) Standardized betas across the cortex by class and network, median-ordered and tested for distribution differences using Wilcoxon's rank-sum test with Bonferroni's correction within class (six comparisons) or network (21 comparisons).

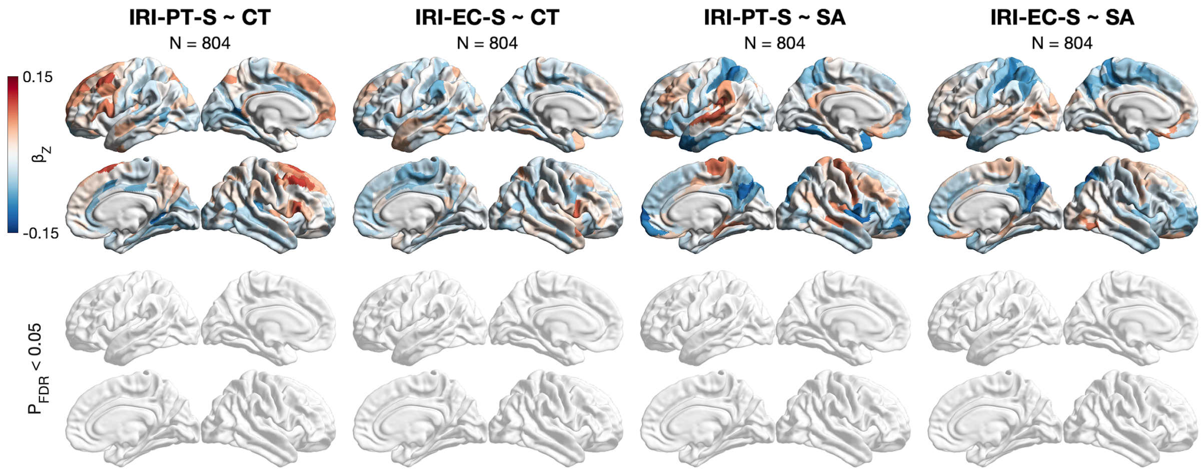


**Figure S4. CT and SA in relation to empathy: Sensitivity analyses.** These versions of IRI-PT and IRI-EC included positively scored items only. All models controlled for age and IQ while the SA models additionally for TIV with FDR correction.

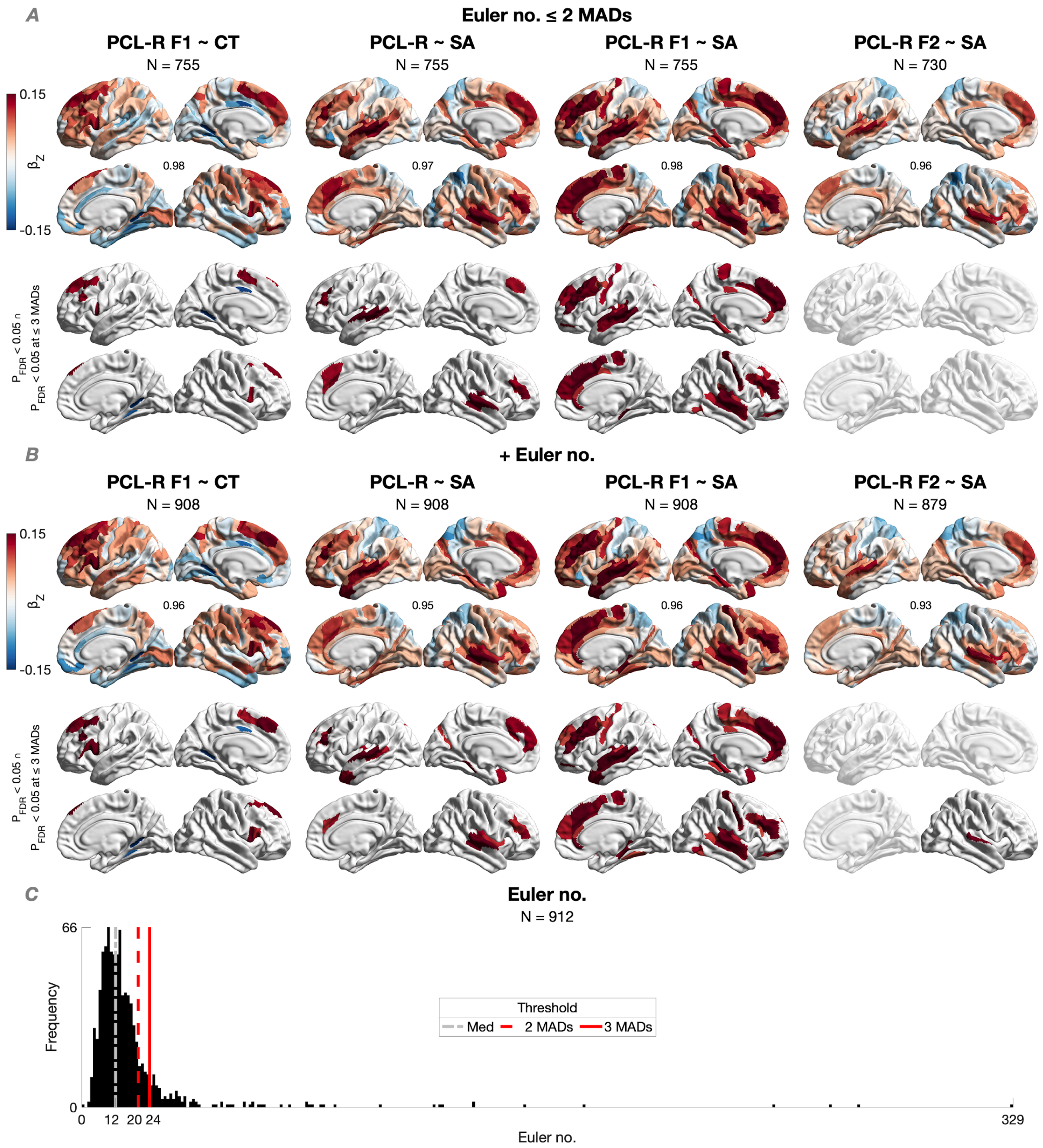


**Figure S5. CT and SA in relation to psychopathy: Sensitivity analyses.** (A) CT and SA in relation to PCL-R total, PCL-R F1, and/or PCL-R F2 following Euler thresholding at a much more conservative threshold (i.e., > 2 median absolute deviations [MADs] above the median, excluding ~17% of participants) compared to the main analysis (i.e., > 3 MADs above the median, excluding ~12% of participants). The FDR-thresholded maps represent the regional intersection at both Euler thresholds (in other words, only standardized betas of the parcels that survived FDR correction at each Euler threshold are plotted). (B) CT and SA in relation to PCL-R total, PCL-R F1, and/or PCL-R F2 where no thresholding was applied, but the Euler number was controlled for as a covariate instead. As in panel A, the FDR-thresholded maps represent the regional intersection with the main analysis. Across the panels, all models controlled for age and IQ, while SA models additionally controlled for TIV. Values in the middle of the unthresholded maps denote Spearman's ρ with the map from the main analysis (all $Ps_{spin} = 0$, $N_{spin} = 1{,}000$). In panel B, sample sizes are N < 912 because participants with missing IQ data or an IQ < 70 (N = 4) were excluded, as in the main analysis. (C) Distribution of the Euler number in the total sample before any data exclusion.

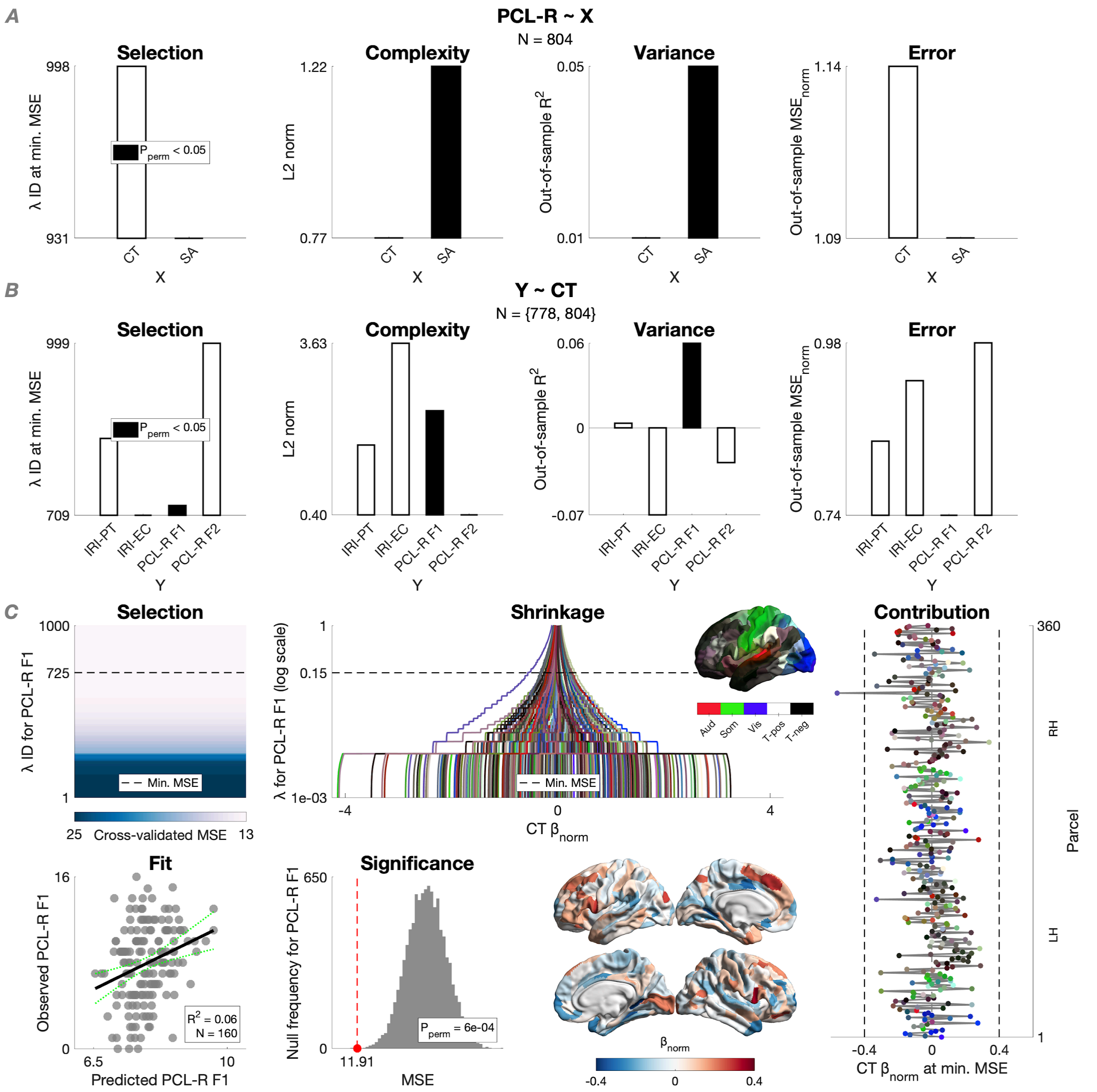


**Figure S6. Multivariate prediction of empathy and psychopathy from CT and/or SA.** (A) For PCL-R total as predicted from CT and SA, we inform on: model selection using cross-validated ridge regression (i.e., lambda corresponding to the minimum cross-validated MSE at which the model was selected); model complexity (i.e., Euclidean norm of the final beta vector); variance explained (i.e., out-of-sample coefficient of determination); and prediction error (i.e., out-of-sample MSE divided by the maximum possible score and thus normalized). CT was corrected for age and IQ while SA additionally for TIV separately in the training (N = 644) and test (N = 160) sets. Only SA was able to predict PCL-R total ($R^2$ = 0.05 [95%: 0.01, 0.07], $P_{perm}$ = 1e-04). (B) For IRI-PT, IRI-EC, PCL-R F1, and PCL-R F2 as predicted from CT, we inform on the same metrics as in panel A. Only PCL-R F1 was able to be predicted ($R^2$ = 0.06 [95% 0.01, 0.10], $P_{perm}$ = 6e-04). (C) For PCL-R F1 as predicted from CT, we inform on model selection, beta shrinkage, final beta vector, predicted-observed fit, and significance based on permutation for out-of-sample MSE ($N_{perm}$ = 10,000).

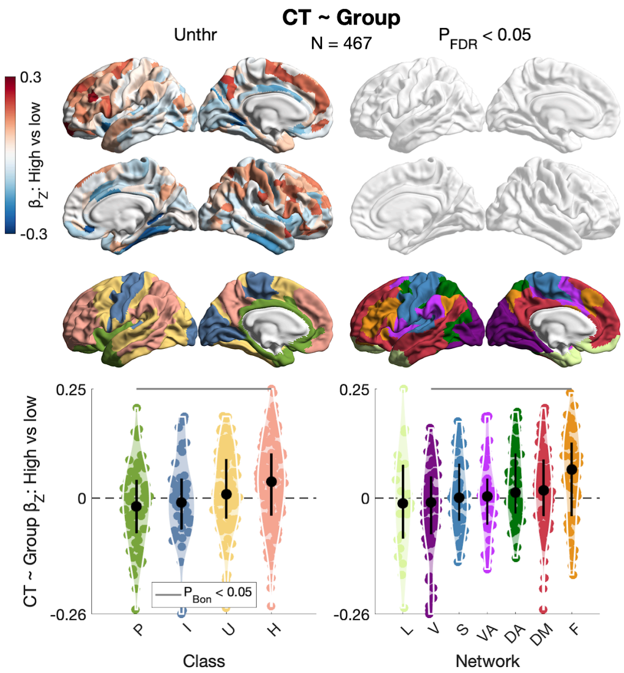


**Figure S7. CT by psychopathy group.** No differences in CT by psychopathy group (high [N = 178] versus low [N = 289]), controlling for age and IQ in a robust linear regression with FDR correction. Standardized betas were median-ordered by class and network, and tested for distribution differences using Wilcoxon's rank-sum test with Bonferroni's correction within class (six comparisons) or network (21 comparisons). Neither was there a difference in mean CT (i.e., globally), controlling for the same covariates ($\beta_Z = 0.08$ [95% CI: -0.10, 0.25], P = 0.390, adj. $R^2 = 0.16$, Cohen's D = 0.04).

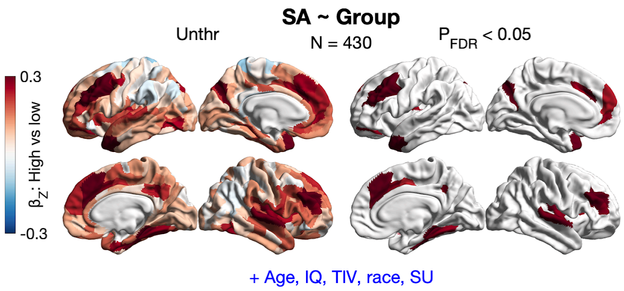


**Figure S8. SA by psychopathy group: Sensitivity analysis.** This model compared high-psychopathy men (N = 159) to low-psychopathy men (N = 271) while controlling for race and total years of substance use in addition to – as in the main analysis (*Fig. **4A***) – age, IQ, and TIV with FDR correction. The unthresholded map correlated at Spearman's $\rho = 0.87$ ($P_{spin}$ = 0, $N_{perm}$ = 1,000) with the main map.

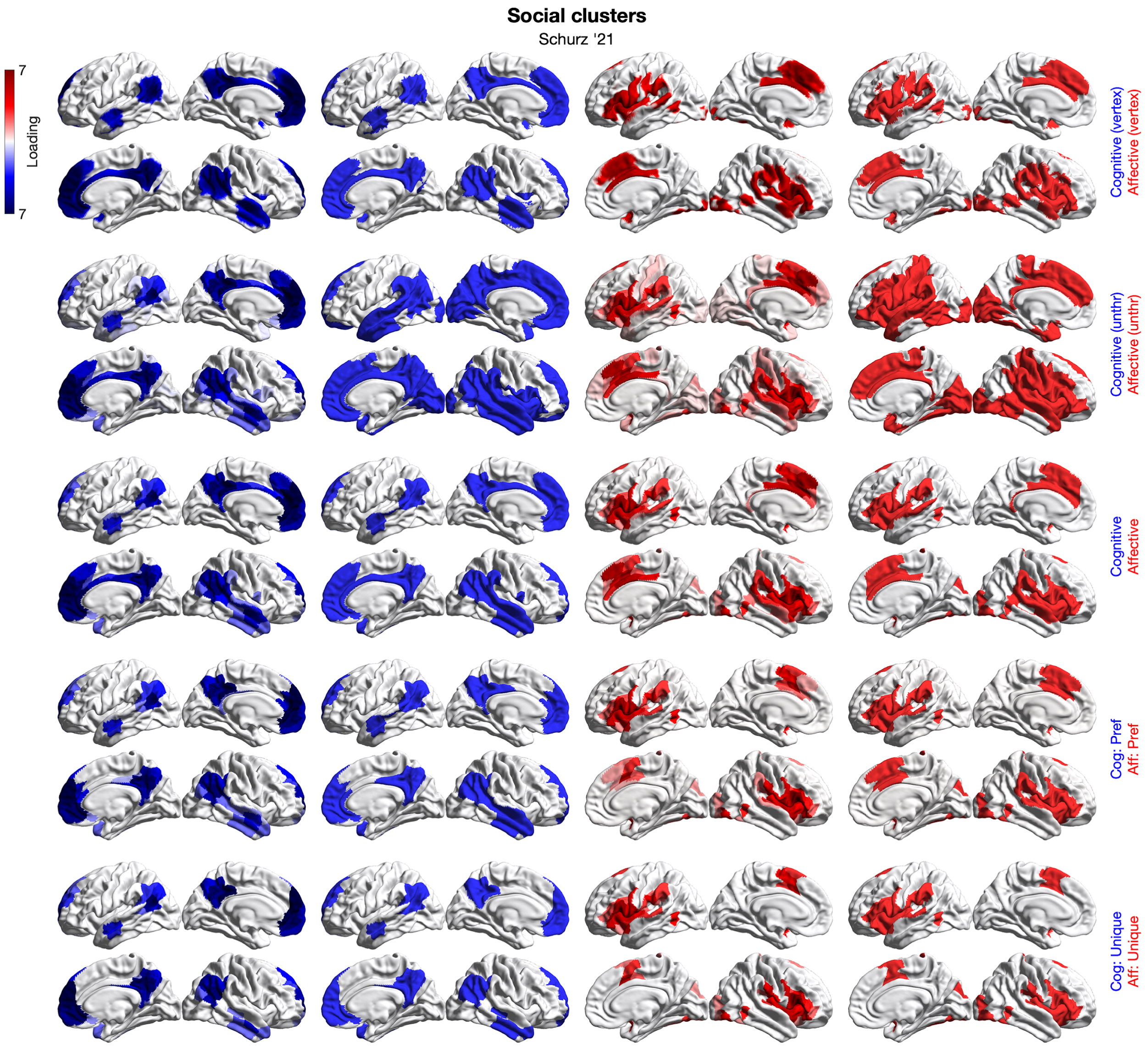


**Figure S9. Meta-analytic clusters of social-cognitive and social-affective processing.** Shown are five pairs of surface-based clusters based on Schurz et al.'s (27) volumetric data across 130 studies. Cognitive clusters in blue are shown in columns 1 and 2, while affective clusters in red are shown in columns 3 and 4 (columns 2 and 4 show cluster masks, i.e., non-zero parcels). Briefly, vertex clusters were parcellated and mean-thresholded ("Cognitive" and "Affective"), subtracted from each other for preferential parcels ("Cog: Pref" and "Aff: Pref"), and masked for unique parcels ("Cog: Unique" and "Aff: Unique").

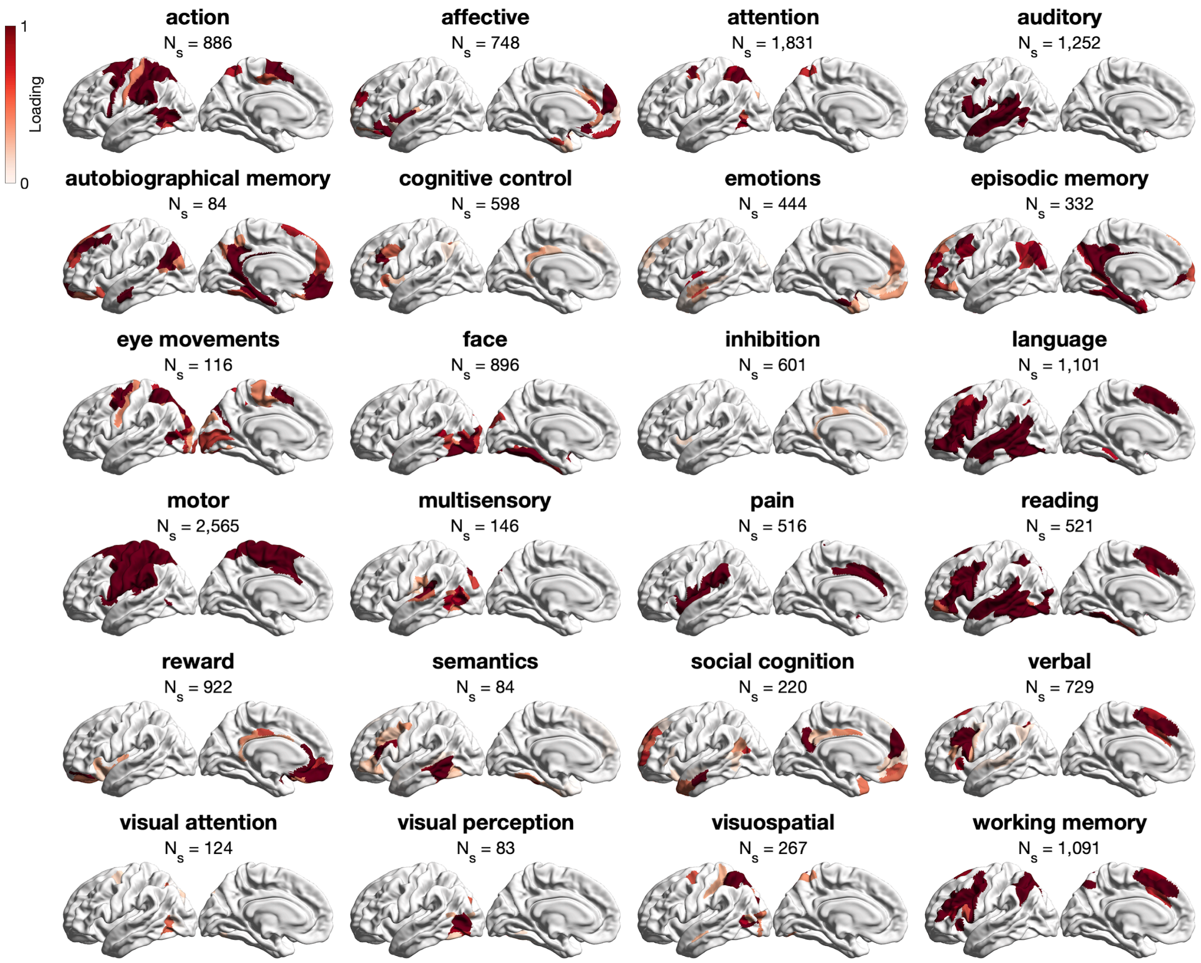


**Figure S10. Neurosynth clusters.** The 24 mean-thresholded Neurosynth clusters whose masks were used to compute the spatial overlap with SA by psychopathy group at $P_{FDR} < 0.05$ in *Fig. **4***. For brevity, we show the left hemisphere only. "$N_s$" = number of studies.

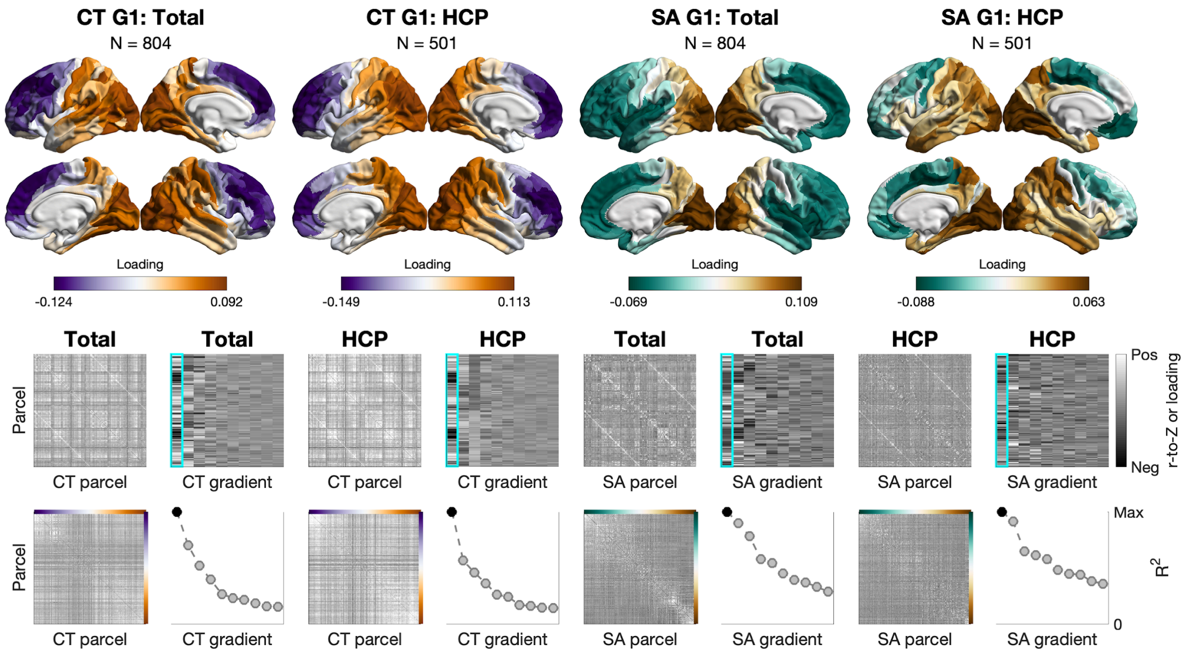


**Figure S11. Macroscale organization of CT and SA in the total and HCP samples.** Shown are raw gradients. Consider the two-by-two left-hand tiles: Sample-specific structural-covariance matrix (top left), array of the first 10 gradients (top right), the matrix ordered by the primary gradient (bottom left), and the first 10 gradients ordered by the proportion of variance explained (i.e., scaled eigenvalues; bottom right). All matrices were set to the range [-0.5, 0.5]; all arrays were set to the minimum-maximum range.

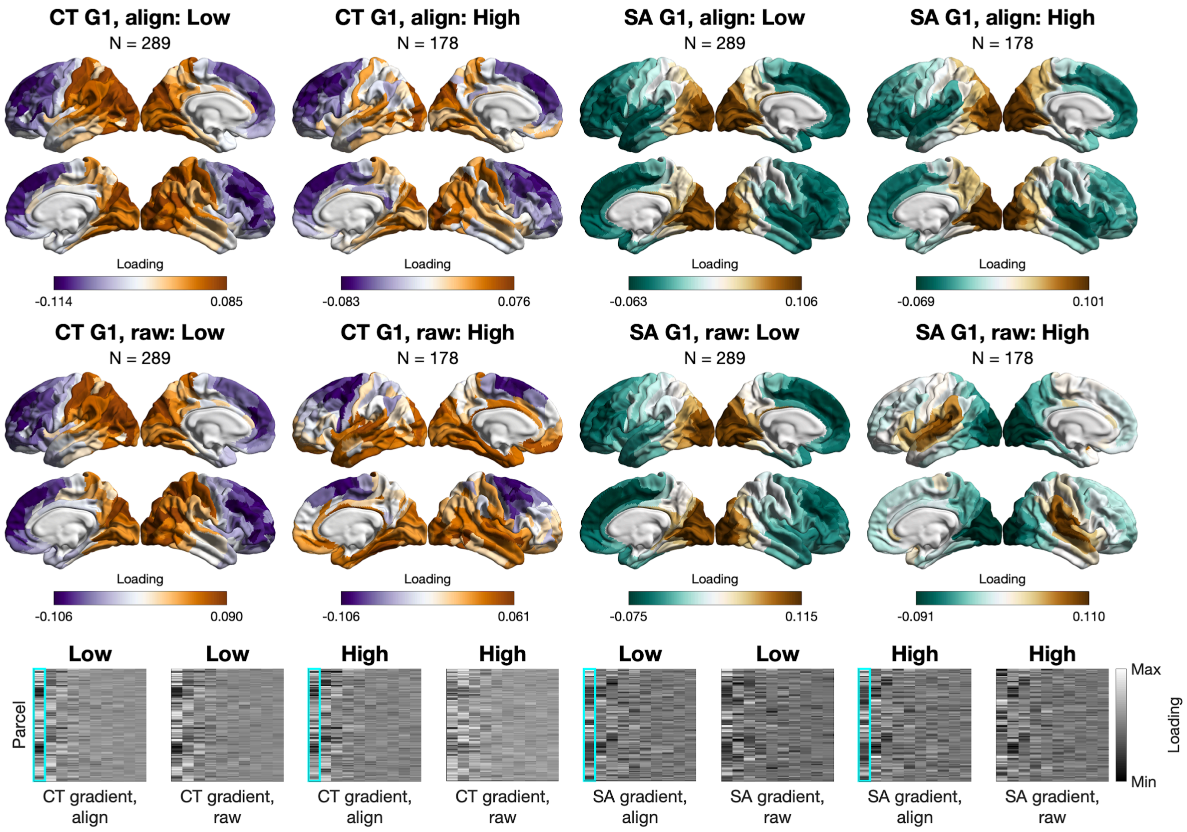


**Figure S12. Macroscale organization of CT and SA by psychopathy group.** Shown are both raw and aligned gradients (aligned via Procrustes rotation to those in the total sample sample, N = 804). At the bottom: Array of the first 10 gradients, both raw and aligned, ordered by the proportion of variance explained (i.e., scaled eigenvalues).

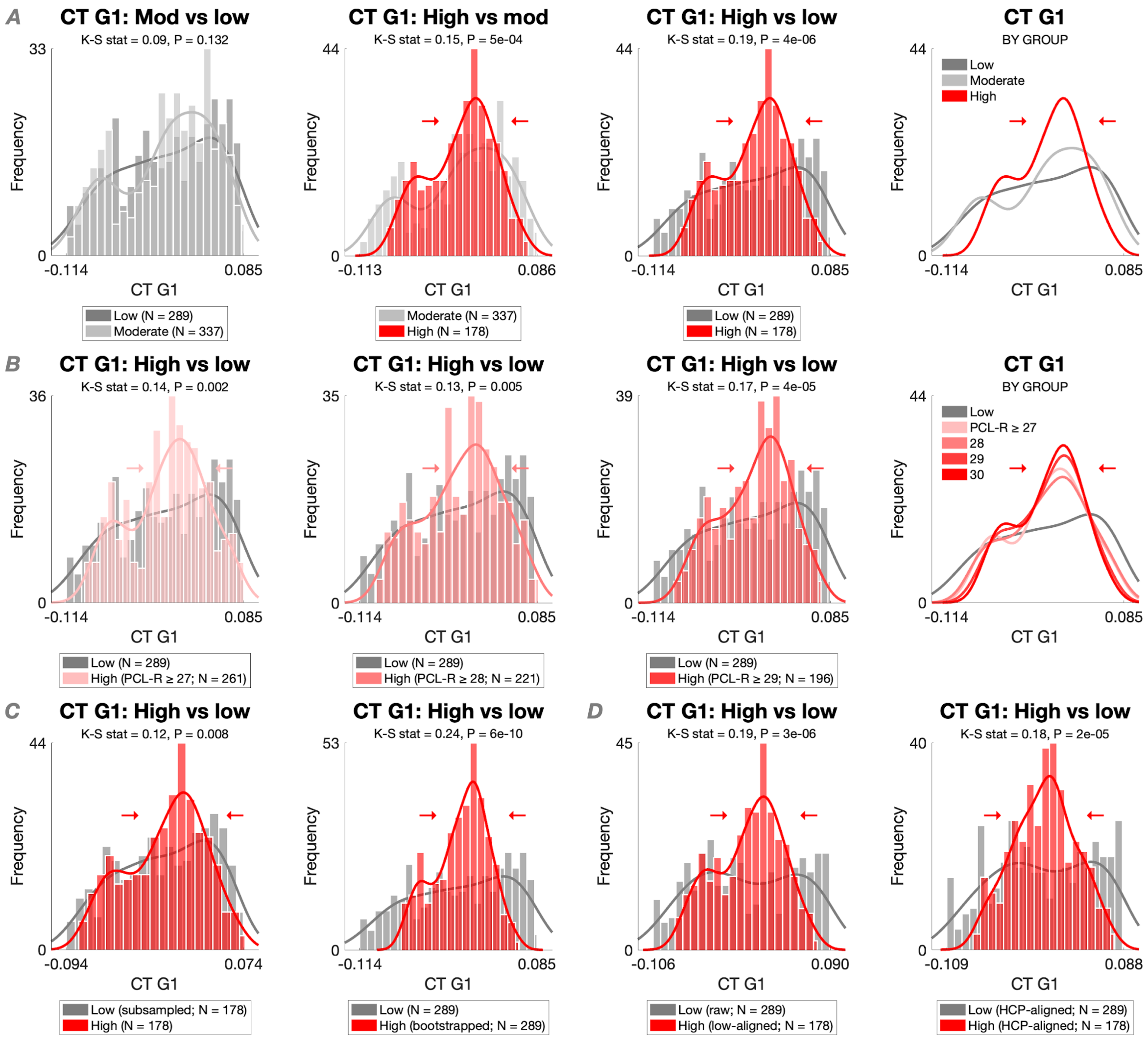


**Figure S13. Macroscale organization of CT by psychopathy group: Global sensitivity analyses.** (A) The CT gradient was compressed in high-psychopathy men (red) compared to both low-psychopathy men (dark gray) and moderate-psychopathy men (to a smaller extent; light gray) using Kolmogorov-Smirnov's test. At the same time, there was no difference between moderate- and low-psychopathy men. (B) Consistently, the CT gradient was compressed in high-psychopathy men compared to low-psychopathy men when using more liberal high-psychopathy thresholds (i.e., PCL-R $\geq$ 27, 28, and 29) instead of the conventional threshold (i.e., PCL-R $\geq$ 30). (C) Consistently, the CT gradient was compressed in high-psychopathy men compared to low-psychopathy men when matching the samples for size: by subsampling the larger low-psychopathy sample ($N_{sub}$ = 1,000; left tile), or by bootstrapping the smaller high-psychopathy sample ($N_{boot}$ = 1,000; right tile). (D) Consistently, the CT gradient was compressed in high-psychopathy men compared to low-psychopathy men when using different templates for Procrustes alignment: (raw) gradients in low-psychopathy men to align (raw) gradients in high-psychopathy men (left tile), or (raw) HCP gradients to align (raw) gradients in both samples (right tile).

**Table S1. Participant characteristics in the HCP sample (N = 501)**

| | | | | | |
|---|---|---|---|---|---|
| N | | 501 | Race (W) | | 385 |
| Age | | 27.91 ± 3.61 | TIV | | 1.71e+06 ± 1.4e+05 |
| | *Range* | [22, 37] | | *Range* | [1.1e+06, 2.1e+06] |
| IQ | | 123.56 ± 14.77 | Euler no. | | 50.41 ± 16.72 |
| | *Range* | [84.55, 153.36] | | *Range* | [11, 116] |

*Note.* Given are means and standard deviations (or frequencies for race) in the male HCP sample. IQ = NIH Toolbox Cognition Total Composite Score: Unadjusted (“CogTotalComp_Unadj”); Race (W) = White; TIV = estimated total intracranial volume [$mm^3$]; Euler no. = total number of topological defects in the cortical surface prior to fixing in the FreeSurfer pipeline.

**Table S2. IRI-PT and IRI-EC items**

| Perspective Taking | Empathic Concern |
|---|---|
| 3.[R] I sometimes find it difficult to see things from the "other guy's" point of view. | 2. I often have tender, concerned feelings for people less fortunate than me. |
| 8. I try to look at everybody's side of a disagreement before I make a decision. | 4.[R] Sometimes I don't feel very sorry for other people when they are having problems. |
| 11. I sometimes try to understand my friends better by imagining how things look from their perspective. | 9. When I see someone being taken advantage of, I feel kind of protective towards them. |
| 15.[R] If I'm sure I'm right about something, I don't waste much time listening to other people's arguments. | 14.[R] Other people's misfortunes do not usually disturb me a great deal. |
| 21. I believe that there are two sides to every question and try to look at them both. | 18.[R] When I see someone being treated unfairly, I sometimes don't feel very much pity for them. |
| 25. When I'm upset at someone, I usually try to "put myself in his shoes" for a while. | 20. I am often quite touched by things that I see happen. |
| 28. Before criticizing somebody, I try to imagine how I would feel if I were in their place. | 22. I would describe myself as a pretty soft-hearted person. |

*Note.* [R] = reverse item.

**Table S3. PCL-R items by factor and facet**

| Factor 1: Interpersonal/Affective | | Factor 2: Lifestyle/Antisocial | |
|---|---|---|---|
| **Facet 1: Interpersonal** | **Facet 2: Affective** | **Facet 3: Lifestyle** | **Facet 4: Antisocial** |
| 1. Glibness/superficial charm | 6. Lack of remorse or guilt | 3. Need of stimulation/proneness to boredom | 10. Poor behavioral control |
| 2. Grandiose sense of self-worth | 7. Emotional shallowness | 9. Parasitic lifestyle | 12. Early behavioral problems |
| 4. Pathological lying | 8. Callousness/lack of empathy | 13. Lack of realistic, long-term goals | 18. Juvenile delinquency |
| 5. Conning/manipulation | 16. Failure to accept responsibility for own actions | 14. Impulsivity | 19. Revocation of conditional release |
| | | 15. Irresponsibility | 20. Criminal versatility |

*Note.* Items 11 (promiscuous sexual behavior) and 17 (many short-term marital relationships) contribute to the total score only.

**Table S4. IRI-PT and IRI-EC by PCL-R variables**

| $\beta_Z$ | [95% CI] | P | $P_{Bon}$ | Adj. $R^2$ | Cohen's D | N |
|---|---|---|---|---|---|---|
| **M1. IRI-PT ~ PCL-R total** (+ age, IQ) | | | | | | |
| -0.111 | [-0.18, -0.04] | 0.002 | 0.004 | 0.026 | – | 804 |
| **M2. IRI-EC ~ PCL-R total** (+ age, IQ) | | | | | | |
| -0.141 | [-0.21, -0.07] | 6e-05 | 1e-04 | 0.052 | – | 804 |
| **M3. IRI-PT ~ PCL-R F1** (+ age, IQ) | | | | | | |
| -0.002 | [-0.07, 0.07] | 0.946 | – | 0.015 | – | 804 |
| **M4. IRI-EC ~ PCL-R F1** (+ age, IQ) | | | | | | |
| -0.108 | [-0.18, -0.04] | 0.002 | 0.004 | 0.045 | – | 804 |
| **M5. IRI-PT ~ PCL-R F2** (+ age, IQ) | | | | | | |
| -0.187 | [-0.26, -0.12] | 3e-07 | 5e-07 | 0.047 | – | 778 |
| **M6. IRI-EC ~ PCL-R F2** (+ age, IQ) | | | | | | |
| -0.135 | [-0.20, -0.06] | 2e-04 | 4e-04 | 0.050 | – | 778 |
| **M7. IRI-PT ~ Psychopathy group** (+ age, IQ) | | | | | | |
| -0.296 | [-0.48, -0.11] | 0.002 | 0.004 | 0.050 | -0.307 | 467 |
| **M8. IRI-EC ~ Psychopathy group** (+ age, IQ) | | | | | | |
| -0.458 | [-0.64, -0.28] | 1e-06 | 2e-06 | 0.085 | -0.503 | 467 |

*Note.* Models are based on robust linear regression. Bonferroni's correction was applied across the IRI subscales. Cohen's D for high- versus low-psychopathy men was computed on raw residuals from the same model but excluding psychopathy group for covariate-corrected estimates. The dependent and continuous independent variables were Z-scored for standardized betas.

**Table S5. IRI-PT and IRI-EC by PCL-R variables: Sensitivity analyses**

| $\beta_Z$ | [95% CI] | P | $P_{Bon}$ | Adj. $R^2$ | Cohen's D | N |
|---|---|---|---|---|---|---|
| **M1. IRI-PT ~ PCL-R total** (+ age, IQ, IRI-EC) | | | | | | |
| -0.039 | [-0.10, 0.02] | 0.214 | – | 0.264 | – | 804 |
| **M2. IRI-EC ~ PCL-R total** (+ age, IQ, IRI-PT) | | | | | | |
| -0.087 | [-0.15, -0.03] | 0.004 | 0.007 | 0.295 | – | 804 |
| **M3. IRI-PT ~ PCL-R F1** (+ age, IQ, IRI-EC) | | | | | | |
| 0.054 | [-0.01, 0.12] | 0.085 | – | 0.264 | – | 804 |
| **M4. IRI-EC ~ PCL-R F1** (+ age, IQ, IRI-PT) | | | | | | |
| -0.103 | [-0.16, -0.04] | 5e-04 | 0.001 | 0.299 | – | 804 |
| **M5. IRI-PT ~ PCL-R F2** (+ age, IQ, IRI-EC) | | | | | | |
| -0.120 | [-0.18, -0.06] | 2e-04 | 4e-04 | 0.272 | – | 778 |
| **M6. IRI-EC ~ PCL-R F2** (+ age, IQ, IRI-PT) | | | | | | |
| -0.051 | [-0.11, 0.01] | 0.102 | – | 0.290 | – | 778 |
| **M7. IRI-PT ~ Psychopathy group** (+ age, IQ, IRI-EC) | | | | | | |
| -0.078 | [-0.24, 0.09] | 0.356 | – | 0.311 | -0.042 | 467 |
| **M8. IRI-EC ~ Psychopathy group** (+ age, IQ, IRI-PT) | | | | | | |
| -0.273 | [-0.42, -0.12] | 5e-04 | 9e-04 | 0.350 | -0.390 | 467 |
| **M9. IRI-PT ~ Psychopathy group** (+ age, IQ, IRI-EC, race, SU) | | | | | | |
| -0.136 | [-0.31, 0.04] | 0.134 | – | 0.332 | -0.106 | 430 |
| **M10. IRI-EC ~ Psychopathy group** (+ age, IQ, IRI-PT, race, SU) | | | | | | |
| -0.295 | [-0.46, -0.13] | 6e-04 | 0.001 | 0.350 | -0.378 | 430 |

*Note.* Models are based on robust linear regression. Bonferroni's correction was applied across the IRI subscales. Cohen's D for high- versus low-psychopathy men was computed on raw residuals from the same model but excluding psychopathy group for covariate-corrected estimates. The dependent and continuous independent variables were Z-scored for standardized betas. SU = total years of substance use based on the Addiction Severity Index.

models (MASEM) of the Interpersonal Reactivity Index (IRI). *Journal of Psychopathology and Behavioral Assessment, 45*(4), 952–963. https://doi.org/10.1007/s10862-023-10098-w

15. Lauterbach, O., & Hosser, D. (2007). Assessing empathy in prisoners--A shortened version of the Interpersonal Reactivity Index. *Swiss Journal of Psychology, 66*(2), 91–101. https://doi.org/10.1024/1421-0185.66.2.91
16. Aharoni, E., & Kiehl, K. A. (2013). Evading justice: Quantifying criminal success in incarcerated psychopathic offenders. *Criminal Justice and Behavior, 40*(6), 629–645. https://doi.org/10.1177/0093854812463565
17. Wechsler, D. (1997). *Wechsler Adult Intelligence Scale (3rd ed.).* Psychological Corporation.
18. Wechsler, D. (2011). *Wechsler Abbreviated Scale of Intelligence (2nd ed.).* Pearson.
19. McLellan, A. T., Kushner, H., Metzger, D., Peters, R., Smith, I., Grissom, G., … Argeriou, M. (1992). The fifth edition of the Addiction Severity Index. *Journal of Substance Abuse Treatment, 9*(3), 199–213. https://doi.org/10.1016/0740-5472(92)90062-S
20. Mesulam, M.-M. (2000). Behavioral neuroanatomy: Large-scale networks, association cortex, frontal syndromes, the limbic system, and hemispheric specializations. In M.-M. Mesulam (Ed.), *Principles of behavioral and cognitive neurology* (2nd ed., pp. 1–120). Oxford University Press.
21. Dorfschmidt, L., Váša, F., White, S. R., Romero-García, R., Kitzbichler, M. G., Alexander-Bloch, A., … Bullmore, E. T. (2024). Human adolescent brain similarity development is different for paralimbic versus neocortical zones. *Proceedings of the National Academy of Sciences of the United States of America, 121*(33), e2314074121. https://doi.org/10.1073/pnas.2314074121
22. Dorfschmidt, L., Váša, F., White, S. R., Romero-García, R., Kitzbichler, M., Alexander-Bloch, A., … Bullmore, E. (2024). Data supporting the manuscript “Human adolescent brain network development is different for paralimbic versus neocortical zones” [Data set]. *Zenodo.* https://doi.org/10.5281/zenodo.11175459
23. Yeo, B. T. T., Krienen, F. M., Sepulcre, J., Sabuncu, M. R., Lashkari, D., Hollinshead, M., … Buckner, R. L. (2011). The organization of the human cerebral cortex estimated by intrinsic functional connectivity. *Journal of Neurophysiology, 106*(3), 1125–1165. https://doi.org/10.1152/jn.00338.2011
24. Hettwer, M. D., Dorfschmidt, L., Puhlmann, L. M. C., Jacob, L. M., Paquola, C., Bethlehem, R. A. I., … Valk, S. L. (2024). Longitudinal variation in resilient psychosocial functioning is associated with ongoing cortical myelination and functional reorganization during adolescence. *Nature Communications, 15*(1), 6283 (2024). https://doi.org/10.1038/s41467-024-50292-2
25. Bzdok, D., Altman, N., & Krzywinski, M. (2018). Statistics versus machine learning. *Nature Methods, 15*(4), 233–234. https://doi.org/10.1038/nmeth.4642
26. Rosenblatt, M., Tejavibulya, L., Jiang, R., Noble, S., & Scheinost, D. (2024). Data leakage inflates prediction performance in connectome-based machine learning models. *Nature Communications, 15*(1), 1829. https://doi.org/10.1038/s41467-024-46150-w
27. Schurz, M., Radua, J., Tholen, M. G., Maliske, L., Margulies, D. S., Mars, R. B., … Kanske, P. (2021). Toward a hierarchical model of social cognition: A neuroimaging meta-analysis and

integrative review of empathy and theory of mind. *Psychological Bulletin, 147*(3), 293–327. https://doi.org/10.1037/bul0000303
28. Schurz, M. (2021). Meta-Analytic Clustering Empathy and ToM. *OSF.* https://doi.org/10.17605/OSF.IO/PAV27
29. Yarkoni, T., Poldrack, R. A., Nichols, T. E., Van Essen, D. C., & Wager, T. D. (2011). Large-scale automated synthesis of human functional neuroimaging data. *Nature Methods, 8*(8), 665–670. https://doi.org/10.1038/nmeth.1635
30. Margulies, D. S., Ghosh, S. S., Goulas, A., Falkiewicz, M., Huntenburg, J. M., Langs, G., … Smallwood, J. (2016). Situating the default-mode network along a principal gradient of macroscale cortical organization. *Proceedings of the National Academy of Sciences of the United States of America, 113*(44), 12574–12579. https://doi.org/10.1073/pnas.1608282113
31. Paquola, C., Vos De Wael, R., Wagstyl, K., Bethlehem, R. A. I., Hong, S.-J., Seidlitz, J., … Bernhardt, B. C. (2019). Microstructural and functional gradients are increasingly dissociated in transmodal cortices. *PLOS Biology, 17*(5), e3000284. https://doi.org/10.1371/journal.pbio.3000284
32. Valk, S. L., Xu, T., Paquola, C., Park, B.-Y., Bethlehem, R. A. I., Vos de Wael, R., … Bernhardt, B. C. (2022). Genetic and phylogenetic uncoupling of structure and function in human transmodal cortex. *Nature Communications, 13*(1), 2341. https://doi.org/10.1038/s41467-022-29886-1
33. Buckner, R. L., Krienen, F. M., Castellanos, A., Diaz, J. C., & Yeo, B. T. T. (2011). The organization of the human cerebellum estimated by intrinsic functional connectivity. *Journal of Neurophysiology, 106*(5), 2322–2345. https://doi.org/10.1152/jn.00339.2011
34. Wu, J., Ngo, G. H., Greve, D., Li, J., He, T., Fischl, B., … Yeo, B. T. T. (2018). Accurate nonlinear mapping between MNI volumetric and FreeSurfer surface coordinate systems. *Human Brain Mapping, 39*(9), 3793–3808. https://doi.org/10.1002/hbm.24213
35. Markello, R. D., Hansen, J. Y., Liu, Z.-Q., Bazinet, V., Shafiei, G., Suárez, L. E., … Misic, B. (2022). neuromaps: Structural and functional interpretation of brain maps. *Nature Methods, 19*(11), 1472–1479 (2022). https://doi.org/10.1038/s41592-022-01625-w
36. Vos de Wael, R., Benkarim, O., Paquola, C., Larivière, S., Royer, J., Tavakol, S., … Bernhardt, B. C. (2020). BrainSpace: A toolbox for the analysis of macroscale gradients in neuroimaging and connectomics datasets. *Communications Biology, 3*(1), 103. https://doi.org/10.1038/s42003-020-0794-7
37. Larivière, S., Park, B-Y., Royer, J., DeKraker, J., Ngo, Sahlas, E., … Bernhardt, B. C. (2024). Connectome reorganization associated with temporal lobe pathology and its surgical resection. *Brain, 147*(7), 2483–2495. https://doi.org/10.1093/brain/awae141
38. Park, B.-Y., Kebets, V., Larivière, S., Hettwer, M. D., Paquola, C., van Rooij, D., … Bernhardt, B. C. (2022) Multiscale neural gradients reflect transdiagnostic effects of major psychiatric conditions on cortical morphology. *Communications Biology, 5*(1), 1024. https://doi.org/10.1038/s42003-022-03963-z
39. Royer, J., Larivière, S., Rodriguez-Cruces, R., Cabalo, D. G., Tavakol, S., Auer, H., … Bernhardt, B. C. (2023). Cortical microstructural gradients capture memory network reorganization in temporal lobe epilepsy. *Brain, 146*(9), 3923–3937. https://doi.org/10.1093/brain/awad125

40. Valk, S. L., Xu, T., Margulies, D. S., Masouleh, S. K., Paquola, C., Goulas, A., … Eickhoff, S. B. (2020). Shaping brain structure: Genetic and phylogenetic axes of macroscale organization of cortical thickness. *Science Advances, 6*(39), eabb3417. https://doi.org/10.1126/sciadv.abb3417
41. Coifman, R. R., Lafon, S., Lee, A. B., Maggioni, M., Nadler, B., Warner, F., & Zucker, S. W. (2005). Geometric diffusions as a tool for harmonic analysis and structure definition of data: Diffusion maps. *Proceedings of the National Academy of Sciences of the United States of America, 102*(21), 7426–7431. https://doi.org/10.1073/pnas.0500334102
42. Mokros, A., Hare, R. D., Neumann, C. S., Santtila, P., Habermeyer, E., & Nitschke, J. (2015). Variants of psychopathy in adult male offenders: A latent profile analysis. *Journal of Abnormal Psychology, 124*(2), 372–386. https://doi.org/10.1037/abn0000042
43. Kim, S., Yoo, S., Xie, K., Royer, J., Larivière, S., Byeon, K., … Park, B.-Y. (2024). Comparison of different group-level templates in gradient-based multimodal connectivity analysis. *Network Neuroscience, 8*(4), 1009–1031. https://doi.org/10.1162/netn_a_00382
44. Burghart, M., & Mier, D. (2022). No feelings for me, no feelings for you: A meta-analysis on alexithymia and empathy in psychopathy. *Personality and Individual Differences, 194*(4), 111658. https://doi.org/10.1016/j.paid.2022.111658
45. Campos, C., Pasion, R., Azeredo, A., Ramião, E., Mazer, P., Macedo, I., Barbosa, F. (2022). Refining the link between psychopathy, antisocial behavior, and empathy: A meta-analytical approach across different conceptual frameworks. *Clinical Psychology Review, 94,* 102145. https://doi.org/10.1016/j.cpr.2022.102145
46. De Brito, S. A., McDonald, D., Camilleri, J. A., & Rogers, J. C. (2021). Cortical and subcortical gray matter volume in psychopathy: A voxel-wise meta-analysis. *Journal of Abnormal Psychology, 130*(6), 627–640. https://doi.org/10.1037/abn0000698
47. Warrier, V., Stauffer, E. M., Huang, Q. Q., Wigdor, E. M., Slob, E. A. W., Seidlitz, J., … Bethlehem, R. A. I. (2023). Genetic insights into human cortical organization and development through genome-wide analyses of 2,347 neuroimaging phenotypes. *Nature Genetics, 55*(9), 1483–1493. https://doi.org/10.1038/s41588-023-01475-y
48. Cohen, J. (1988). *Statistical power analysis for the behavioral sciences (2nd ed.).* Lawrence Erlbaum Associates.
